# Plasmid transmitted small multidrug resistant (SMR) efflux pumps differ in gene regulation and enhance tolerance to quaternary ammonium compounds (QAC) when grown as biofilms

**DOI:** 10.1101/768630

**Authors:** Carmine J. Slipski, Taylor R. Jamieson, Amy Lam, Vanessa Leung Shing, Kelli Bell, George G. Zhanel, Denice C. Bay

## Abstract

Small multidrug resistance (SMR) efflux pump genes are commonly identified from integrons carried by multidrug-resistant (MDR) plasmids. SMR pumps are annotated as ‘*qac*’ for their ability to confer resistance to quaternary ammonium compounds (QACs) but few *qac* are characterized to date. Hence, we have examined SMR sequence diversity, antimicrobial susceptibility, and gene expression from >500 sequenced proteobacterial plasmids. SMR sequence diversity from plasmid database surveys identified 20 unique SMR sequences annotated as *qacE/EΔ1/F/G/H/I/L*, or *sugE*. Phylogenetic analysis shows ‘Qac’ sequences are homologous to archetypical SMR member EmrE, and share a single sequence origin. In contrast, SugE sequences are homologous to archetypical member Gdx/SugE and likely originate from different species. SMR genes, *qacE, qacEΔ1, qacF, qacG, qacH*, and *sugE(p)*, were over-expressed in *Escherichia coli* to determine their QAC antimicrobial susceptibility as planktonic, colony, and biofilms. SMRs (except *qacEΔ1/sugE*) expressed in biofilms significantly increased its QAC tolerance as compared to planktonic and colony growth. Analysis of upstream SMR nucleotide regions indicate *sugE(p)* genes are regulated by type II guanidinium riboswitches, whereas *qacE* and *qacEΔ1* have a conserved class I integron Pq promoter, and *qacF/G/H* are regulated by integron Pc promoter in variable cassettes region. Beta-galactosidase assays were used to characterize growth conditions regulating Pq and Pc promoters and revealed that Pq and Pc have different expression profiles during heat, peroxide, and QAC exposure. Altogether, this study reveals that biofilm growth methods are optimal for SMR-mediated QAC susceptibility testing and suggests SMR gene regulation on plasmids is similar to chromosomally inherited SMR members.

## Introduction

Small multidrug resistance (SMR) proteins belong to a family of proton motive force driven, multidrug selective efflux pumps found in bacteria (and archaea) that are small (100-150 amino acids) and composed of 4 transmembrane (TM) α-helices as compared to most efflux pump family members with 12-14 TM α-helices (1). Depending on their phylogeny, SMR members confer low to moderate levels (2-8 fold increases in in minimum inhibitory concentration (MIC) values from controls) of tolerance (reduced susceptibility or resistance) to a variety of guanidinium (Gdm^+^) and lipophilic cation containing chemical compounds when they are overexpressed in bacteria (2–5). The most notable Gdm^+^ compounds associated with SMRs are quaternary ammonium compounds (QACs). QACs include a wide range of disinfectants and antiseptics (e.g. benzalkonium and cetrimide) as well as DNA/RNA inter-chelating lipophilic dyes (e.g. ethidium, acriflavine) (6–8). SMR family members that confer antimicrobial tolerance by the expression of a single gene copy can be divided into two major groups, 1) small multidrug proteins (SMPs) and 2) suppressor of *groEL* mutations (SUG) subclass, recently renamed as the ‘riboswitch regulated Gdm^+^ exporter’ (GDM) subclass. GDM were formerly named “suppressor of the *groEL* mutations” based on an incorrectly identified cloning artifact (4, 9). One of the most well characterized GDM subclass members is *E. coli* Gdx (formerly SugE), which confers tolerance to small molecular weight Gdm^+^ compounds (2) and/or to a limited range of QAC antiseptic detergents, such as cetyltrimethylammonium (CTAB) and cetylpyridinium (CTP) (4, 9–11). GDM members from various taxa are regulated by different Gdm^+^ binding riboswitches (types I, II, and III) and phylogenetically associate based on riboswitch types (2). Riboswitch sequences are located in the 5’ untranslated region (UTR) of the *gdx/sugE* transcript, where the UTR adopts stem loops arranged in an “off” conformation, arresting protein translation. Translation is only switched “on” when a Gdm^+^ containing molecule binds to the UTR stem loop permitting translation of the RNA (as reviewed by (12)). In contrast to GDM, SMP subclass members are typified by the most well characterized SMR member, *E. coli* ethidium multidrug resistance protein E (*emrE*). Unlike GDM proteins, SMP members confer tolerance to a wide range of chemically diverse QACs and cation based antimicrobials including some aminoglycosides (as reviewed by (3, 4, 9)). To date, SMP member regulation by a Gdm^+^ riboswitch has not been demonstrated (2, 4, 9–11), and many SMP members are transmitted by integrons, prophages, and multidrug resistant (MDR) plasmids (2, 9).

In bacteria, efflux is one of the most well studied QAC tolerance mechanisms (13, 14) where SMR members are believed to play an important role in lateral gene transmitted QAC tolerant phenotypes. SMR members are frequently encoded by horizontally transmitted genetic elements, primarily class I integrons mobilized on transposons and MDR plasmids. Integron associated SMRs were annotated to as ‘qac’ genes (*qacC, qacE, qacE*Δ*1, qacF, qacG, qacH, qacI, qacL*) or ‘*sugE*’ genes based on their substrate selectivity or on their SMR sequence homology. ‘*qac*’ genes are considered to be a hallmark of class I integrons as they are often located within the 3’ conserved region of these elements, to lesser extent located within the variable cassette region (13, 15, 16), or mobilized on various MDR plasmids by transposons (17, 18). SMR genes carried by MDR plasmids and integrons, are often monitored in food/ retail meat surveillance facilities and within clinical isolate collections to predict or correlate the extent of bacterial antiseptic/disinfection susceptibility/tolerance (19–24). Most surveillance studies infer antiseptic tolerance from SMR gene presence or absence, despite studies and findings downplaying SMR’s role and contribution to overall antiseptic/ disinfectant tolerance (14, 25). To date, there are no Clinical and Laboratory Standards Institute (CLSI) guidelines for determining QAC or other antiseptics/ disinfectant susceptibility/resistance breakpoints. As a result, we cannot definitively state resistance values for QACs and other biocides using antimicrobial susceptibility testing (AST) methods (as reviewed by (26, 27)); therefore, we use the recommended term ‘tolerance’ in lieu of ‘resistance’ for biocides including QACs. Very few Gram-negative plasmid encoded SMR members (*qacE, qacE*Δ*1*, and *qacF*) have been directly cloned and over-expressed for AST, and these methods generally involve broth dilution or agar spotting methods (13, 28–30). Most plasmid encoded SMR members, *qacG, qacH, qacI, qacL* and *sugE* have not been experimentally characterized to date, highlighting an important knowledge gap to be addressed in SMR studies.

Another knowledge gap concerning SMR efflux pumps, is how differences in SMR-mediated antiseptic tolerance may be influenced by AST growth method(s), i.e. planktonic, colony, or biofilm (26, 31, 32). QAC-based disinfectants solutions are often used to eradicate bacterial biofilms (33, 34) yet, biofilm tolerance to QACs is rarely examined by AST methods. Biofilms represent a physiological growth state that causes planktonic/motile bacteria to attach to solid surfaces and form a sessile surface attached community that is more tolerant to antimicrobials by the secretion of extrapolymeric substances, where cells become less metabolically active within the biofilm. Biofilm growth is common in environmental samples and in chronic animal/human infections (as reviewed by (35, 36)). Enhanced QAC tolerance by bacteria growing as a biofilm is an important concern, since QACs and other biocides are heavily relied upon by food, clinical, agricultural industries to disinfect difficult to sterilize surfaces (as reviewed by (14)). Hence, reductions in QAC biofilm eradication effectiveness result in higher costs for human/animal health and food product safety.

The aim of this study was to perform an in-depth analysis of SMR sequence diversity on MDR plasmids, to identify frequently transmitted SMR sequences, compare SMR member antimicrobial selectivity using AST with bacteria grown as planktonic, colony and biofilms, and explore SMR gene expression/regulation on MDR plasmids. To begin, we bioinformatically surveyed Gram-negative bacterial plasmid sequence databases for SMR proteins to determine the extent of SMR sequence variation, distribution, and diversity. We established SMR phylogenetic relatedness to well characterized SMR subclass members EmrE and Gdx. A total of 20 unique SMR protein sequences were identified from a bioinformatic survey of over 500 SMR encoding plasmids, where six representative SMR sequences were selected (*qacE, qacE*Δ*1, qacF, qacG*, and *qacH*, and *sugE(p)*) for further experimental AST analysis. Six SMR genes were cloned into the expression vector pMS119EH and transformed into *Escherichia coli* K-12 strains BW25113 and KAM32 for AST with a library of 13 antimicrobials (9 QACs, 2 interchelating dyes, and 2 antibiotics). To determine if growth influenced SMR QAC tolerance phenotypes, 3 different AST methods were examined in this study: agar dilution, broth microdilution, and minimal biofilm eradication devices (Innovotech MBEC assays; formerly known as the Calgary biofilm device) representing colony, planktonic and biofilm culture growth physiologies, respectively. All three AST growth methods reconfirmed the current hypothesis that *qac* and *sugE* genes confer distinct antimicrobial substrate selection profiles based on their phylogenetic relationships to either the SMP or GDM subclass of the SMR family. A comparison of AST growth methods tested our second study hypothesis, that SMR over-expression in bacteria (*E. coli*) growing as colony or a biofilm exhibit greater and more significant QAC tolerance as compared to planktonic growth methods. The third hypothesis we examined was that is plasmid/ integron encoded SMRs related to the GDM subclass will be regulated by a Gdm^+^ riboswitch as compared to SMP homologs by a discrete plasmid encoded promoter. Previous studies indicated that *qacEΔ1* genes rely on a ‘Pc’ or a ‘Pq/ Pqac’ promoter region associated with class I integrons (15, 37) but is it unclear how other SMR genes are regulated. To address the third hypothesis, we bioinformatically analyzed the 500 nt upstream region corresponding to each SMR sequence we surveyed to identify if class I integron promoters ‘Pc’ and ‘Pq/ Pqac/ P3’ (as described by (15, 38)) or Gdm^+^ binding riboswitches (as reviewed by (12)) sequences were present on plasmids. Since very little is known about Pq promoter induction, our final aim delved deeper into class I integron ‘Pc’ and ‘Pq’ promoter regulation to determine if exposure to stress, specifically, sub-inhibitory QAC concentrations, heat (42°C), and oxidative (H_2_O_2_) stress, had a similar influence on these promoters identified upstream from specific *qac* genes. By cloning the Pc and Pq promoter regions into a β-galactosidase *lacZ* reporter plasmid we compared β-galactosidase activity using a 96-well microplate assay in *E. coli* promoter transformant that were grown under various growth modes (planktonic and biofilm) and different stress conditions, highlighting Pq and Pc promoter differences. Altogether this study, clarifies three major knowledge gaps related to plasmid/ integron transmitted SMR antimicrobial resistance profiles and offers greater insight into how plasmid encoded SMR gene expression during bacterial growth influences tolerance to a wide range of QACs.

## 2. Results

### 2.1 Bioinformatic analysis of plasmid encoded SMR sequences indicate a relationship to GDM or SMP subclass members

The majority (87-92%) of the plasmids encoding for either a ‘Qac’ or ‘SugE’ sequence in our database surveys were identified from γ-Proteobacteria, specifically the order Enterobacteriales, and to a lesser extent sequence identified from orders Pseudomonadales, Vibrionales, Aeromonadales (Fig. 1B; Table S1). Due to the predominance of SMR genes isolated from Enterobacterial plasmids, no obvious trends were apparent to support a link between specific environmental niches and SMR co-selection. Due to animal and human Enterobacteriales contamination, these species can be detected from and thrive in a wide range of conditions, explaining their detection in diverse environments. Plasmid sequencing biases favoring Enterobacterial clinical isolates as well as food/ retail meat contaminants likely bias SMR predominance within Enterobacteriales (Fig. 1C). Therefore, it is unclear if specific SMR members are preferentially enriched or selected by plasmids/ integrons in specific environmental sources. Regardless, the diversity and environmental distribution of ‘Qac’ and ‘Gdx/ SugE’ from our plasmid database sequence surveys remains consistent with SMR gene detection rates and trends reported by surveillance studies of *E. coli* (20, 22, 24), *Salmonella enterica* spp. (39, 40), *Pseudomonas spp*. (41–43), *Vibrio* spp. (29, 44), and *Acinetobacter* spp. (45, 46).

**Figure 1.**
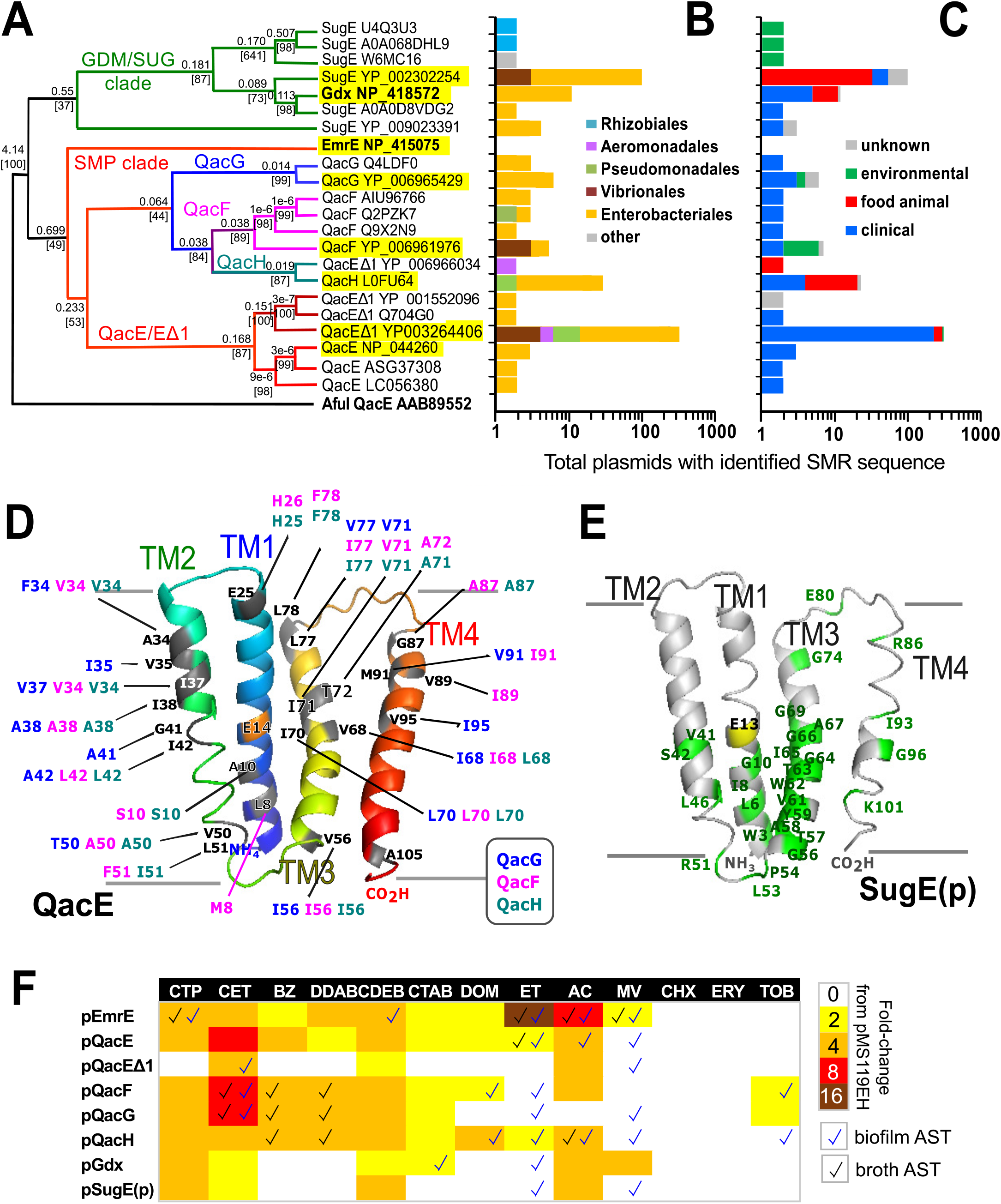
The phylogenetic relatedness, taxonomic distribution, isolation source, sequence variation, AST summary of plasmid encoded SMR members. **A)** A Maximum likelihood (PhyML) dendrogram of the 20 unique Qac and Gdx/ SugE protein sequences identified from Gram-negative plasmid surveys. Branch node confidence values were determined by performing 100 bootstrap replicates; branch distance values are provided above bootstrap confidence values are provided in brackets. Archetypical *E. coli* SMR members, EmrE (NP_415075) and Gdx (NP_044260) as well as the outgroup Archaeal SMR family *Archaeoglobus fulgidus* QacE (AAB89552) sequences are highlighted in bold. Yellow highlighted sequences were selected for AST based on their detection frequency in various plasmids. **B)** The taxonomic origin of SMR containing plasmids corresponding to each sequence listed in panel A based on data in Table S1. **C)** The reported isolation sources of plasmids with verified SMR sequences (summarized from data provided in Table S1). **D)** An I-TASSER (83) homology model of the QacE protein (NP_044260) monomer, shown as a ribbon secondary structure diagram, where amino acid sequence variations (grey) identified in QacF, QacG and QacH alignments (Fig. S2) are highlighted and colour coded according to the inset panel legend. All 4 predicted transmembrane helices (TM) are shown and colour coded (TM1; green, TM2; blue; TM3; yellow; TM4; red). **E)** An I-TASSER homology model of SugE(p) protein (YP_002302254) monomer shown as a grey ribbon secondary structure diagram, where conserved amino acids (green) are highlighted from the sequence alignment shown in (Fig. S2). **F)** A heatmap summary of mean MIC values of KAM32 SMR transformant AST agar spot plate method from Table 3. Check marks on the specific heatmap boxes indicate significant (4-fold increase from pMS119EH control) MIC values were identified in broth AST (black checkmark) or (4-fold increased from control) MBEC biofilm AST (blue checkmark) experiments for the same transformant and antimicrobial combination tested.

SMR protein sequences frequently annotated as ‘Qac’ formed three closely related subclades: i) QacE/QacEΔ1 cluster, ii) QacG, and iii) the QacF/I/L/H clusters (Fig. 1A), where each Qac was distantly related to *E. coli* EmrE (Fig. 1A). QacEΔ1 was the most frequently detected SMP member within diverse environmental and clinical isolates, in agreement with previous SMR surveillance studies (31, 43, 46, 47). This is not surprising, considering QacEΔ1 has high conservation within the 3’ conserved region of Class I integrons, which we identified in 99.5% of plasmid sequences surveyed herein (Table S1, Table S2). Many Qac sequences we examined were misannotated; QacEΔ1 was most frequently misannotated as ‘EmrE’, ‘QacE’, and ‘Smr’, while ‘QacF’ and ‘QacH’ sequences were misannotated as ‘QacE’, ‘QacEΔ1’, ‘QacI’, and ‘QacL’ (Table S1, Table S2-3) and highlights caution when determining SMR sequence association by annotation only. Multiple sequence alignments of all Qac proteins showed high overall sequence identity (66-99% pairwise identities) to each other but not to EmrE suggesting Qac sequences share a close and common sequence origin (Fig. S1). QacEΔ1 had the lowest sequence similarity, due to the an in-frame replacement of the 4^th^ TM strand extending the original QacE protein to 115 aa (Fig. S1) (28). QacE, QacF, QacG, and QacH proteins had the least amino acid variations (≤15 residue differences total) suggesting they are all isoforms with low sequence similarity to EmrE (42-49% pairwise sequence identity; Fig. S1). Most amino acid differences identified between different Qac sequences altered hydrophobic residues located on the lipid-facing surfaces of each TM protein and away from the proposed H^+^ and drug binding active site residue ‘E14’ as shown in the monomeric QacE homology model generated from *E.coli* EmrE X-ray diffraction crystal structure (3B5D.pdb (48); Fig. 1D). This suggests that most sequence variations among QacE, QacF, QacG, and QacH isoforms are less likely to impact substrate recognition and substrate efflux. Altogether, our analysis suggests that plasmid-transmitted ‘qac’ gene sequences in class I integrons have likely originated from a single *qac* origin, and show close relationship to SMP members, in agreement with previous SMR phylogenetic analyses (16, 49).

Phylogenetic analysis of plasmid-encoded GDM members formed two distinct subclades, i) a γ-proteobacterial clade with homology to *E. coli* Gdx (NP_418572) and ii) an α-proteobacterial Rhizobiales ‘SugE’ clade (Fig. 1A, S2). The most frequently identified GDM member was SugE(p) (YP_002302254), which had 83% sequence identity to *E. coli* Gdx and was identified from γ-proteobacterial plasmids isolated from Enterobacteriales, specifically *E. coli* and *Salmonella enterica* species (Table S1; TableS2). SugE(p) was frequently detected on plasmids isolated from species found in contaminated foods and retail animal (primarily poultry) sources (Table S1), suggesting the food industry may select for SugE(p) maintenance on plasmids from these species. Interestingly, 7 GDM sequences we surveyed were 100% identical to chromosomally encoded *E. coli* Gdx (Fig. 1B), indicating that the archetypical *E. coli gdx* selected for this study is also horizontally transmissible on Enterobacterial plasmids. In contrast to plasmid encoded SMP members, all 7 GDM protein pairwise sequence identities were much lower when compared to the archetypical *E.coli* Gdx (25-30%) sequence (Fig. 1E; Fig. S2). Altogether, this suggests that *gdx/ sugE* on plasmids are more likely to be acquired from different sequence origins or species that acquired the plasmid. Alternatively, plasmid GDM sequences may have lower selective pressure exerted to maintain their sequence identity on plasmids, however, horizontal gene transfer makes determination of this difficult to discern using the current plasmid dataset.

### 2.2 AST of E. coli SMR gene transformants show significant QAC tolerance when grown as biofilms in a wildtype efflux pump strain

Six of the most frequently identified plasmid-encoded Qac and Gdx/ SugE(p) sequences we identified from our proteobacterial plasmid surveys, highlighted in yellow in Figure 1A and listed in Table 1, were selected for experimental AST. Each sequence was synthesized, cloned, and overexpressed (by 0.05 mM isopropyl β-D-1-thiogalactopyranoside (IPTG) final concentration) from a low copy number vector, pMS119EH (50), previously used to examine archetypical *E. coli* SMR members *emrE* and *gdx* genes (3, 4). EmrE and Gdx were included in the analysis as comparative SMR subclass member controls. AST analyses were used to calculate the minimal inhibitory concentration (MIC) and minimal biofilm eradication concentration (MBEC) values for vector transformants under three different growth conditions (broth, agar spot colony, and pegged lid biofilms) in two different strains of *E. coli* K-12. *E. coli* BW25113 was selected as a ‘wildtype’ strain (51) and *E. coli* KAM32 (52) was selected as a multiple efflux pump gene (Δ*acrB*, Δ*mdtK*) deletion strain, which is frequently used for efflux pump AST analyses (for an example refer to (53)). AST of SMR vector transformants BW25113 and KAM32 strains respectively were examined with a library of 13 previous tested QACs and antibiotics selected on the basis of their ability to confer antimicrobial tolerance in other SMR over-expression studies (as reviewed by (4)) and results are summarized on Tables 2–3.

**Table 1.**
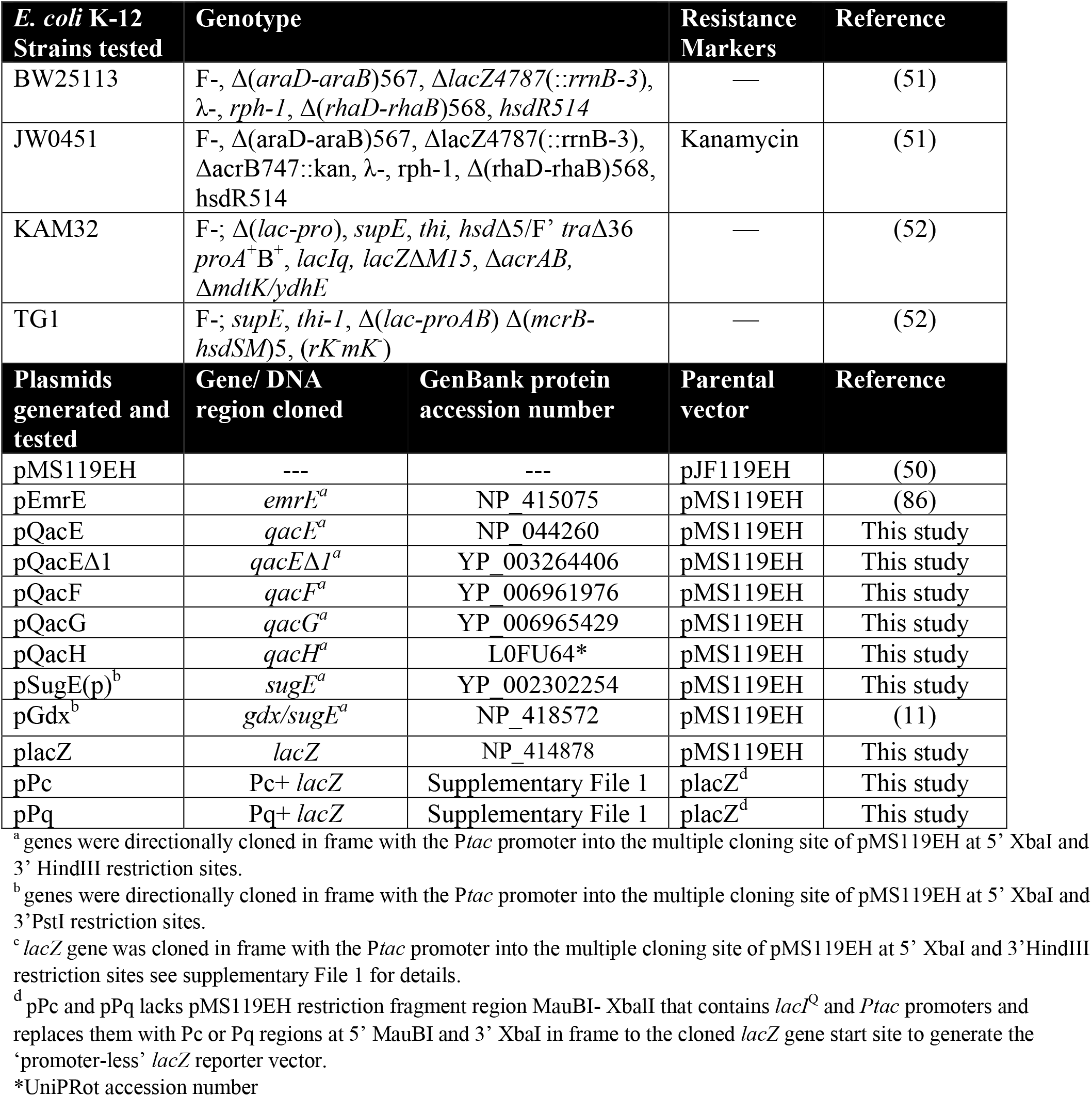
Strains and plasmids used or generated in this study.

**Table 2.**
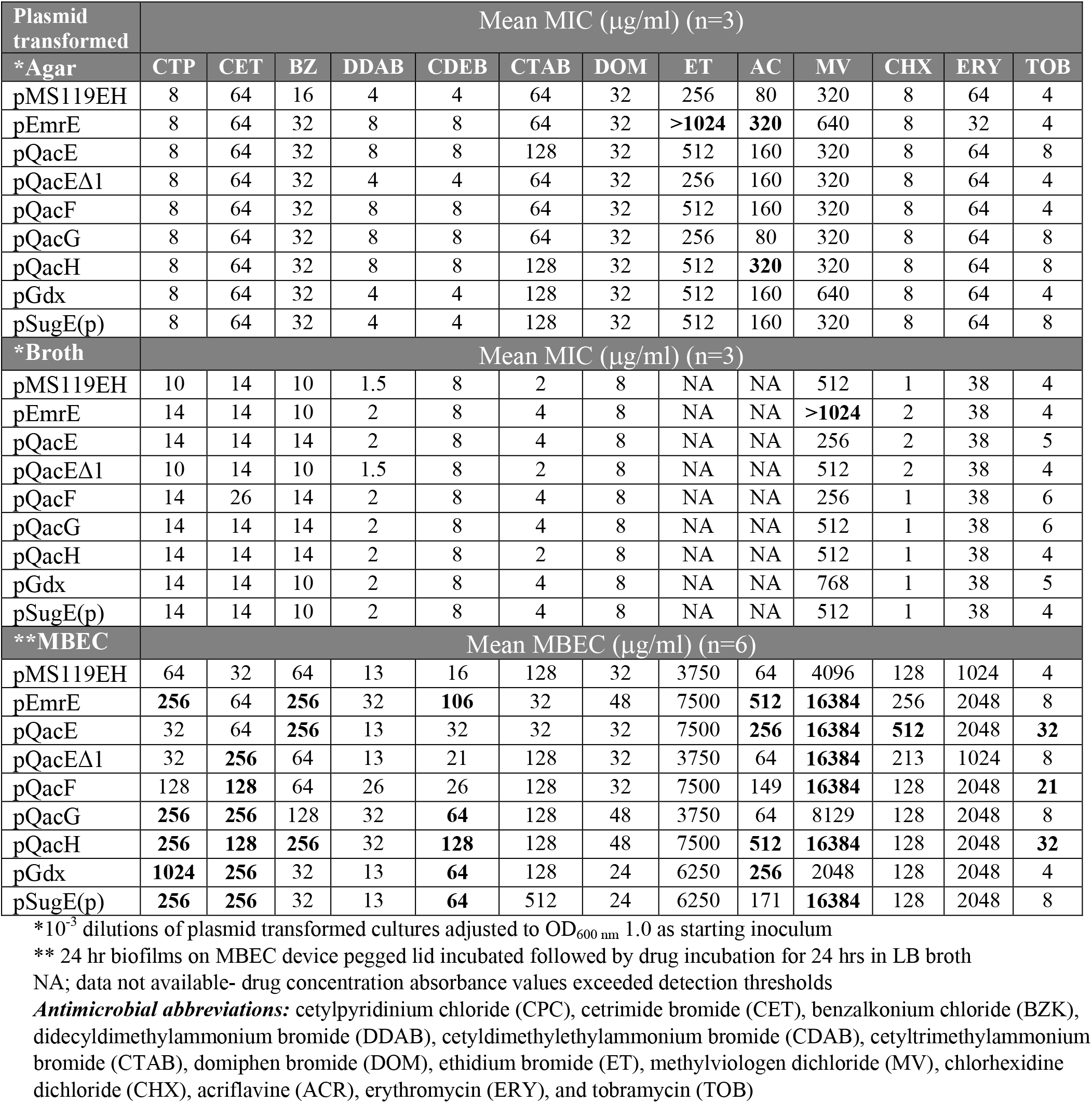
Summary of antimicrobial MIC and MBEC values determined for *E. coli* BW25113 transformed with plasmids expressing various SMR genes grown as planktonic (broth), colony (agar) and biofilms (MBEC).

| Plasmid transformed | Mean MIC (µg/ml) (n=3) |  |  |  |  |  |  |  |  |  |  |  |  |
| --- | --- | --- | --- | --- | --- | --- | --- | --- | --- | --- | --- | --- | --- |
| *Agar | CTP | CET | BZ | DDAB | CDEB | CTAB | DOM | ET | AC | MV | CHX | ERY | TOB |
| pMS119EH | 8 | 64 | 16 | 4 | 4 | 64 | 32 | 256 | 80 | 320 | 8 | 64 | 4 |
| pEmrE | 8 | 64 | 32 | 8 | 8 | 64 | 32 | >1024 | 320 | 640 | 8 | 32 | 4 |
| pQacE | 8 | 64 | 32 | 8 | 8 | 128 | 32 | 512 | 160 | 320 | 8 | 64 | 8 |
| pQacEΔ1 | 8 | 64 | 32 | 4 | 4 | 64 | 32 | 256 | 160 | 320 | 8 | 64 | 4 |
| pQacF | 8 | 64 | 32 | 8 | 8 | 64 | 32 | 512 | 160 | 320 | 8 | 64 | 4 |
| pQacG | 8 | 64 | 32 | 8 | 8 | 64 | 32 | 256 | 80 | 320 | 8 | 64 | 8 |
| pQacH | 8 | 64 | 32 | 8 | 8 | 128 | 32 | 512 | 320 | 320 | 8 | 64 | 8 |
| pGdx | 8 | 64 | 32 | 4 | 4 | 128 | 32 | 512 | 160 | 640 | 8 | 64 | 4 |
| pSugE(p) | 8 | 64 | 32 | 4 | 4 | 128 | 32 | 512 | 160 | 320 | 8 | 64 | 8 |
| *Broth | Mean MIC (µg/ml) (n=3) |  |  |  |  |  |  |  |  |  |  |  |  |
| pMS119EH | 10 | 14 | 10 | 1.5 | 8 | 2 | 8 | NA | NA | 512 | 1 | 38 | 4 |
| pEmrE | 14 | 14 | 10 | 2 | 8 | 4 | 8 | NA | NA | >1024 | 2 | 38 | 4 |
| pQacE | 14 | 14 | 14 | 2 | 8 | 4 | 8 | NA | NA | 256 | 2 | 38 | 5 |
| pQacEΔ1 | 10 | 14 | 10 | 1.5 | 8 | 2 | 8 | NA | NA | 512 | 2 | 38 | 4 |
| pQacF | 14 | 26 | 14 | 2 | 8 | 4 | 8 | NA | NA | 256 | 1 | 38 | 6 |
| pQacG | 14 | 14 | 14 | 2 | 8 | 4 | 8 | NA | NA | 512 | 1 | 38 | 6 |
| pQacH | 14 | 14 | 14 | 2 | 8 | 2 | 8 | NA | NA | 512 | 1 | 38 | 4 |
| pGdx | 14 | 14 | 10 | 2 | 8 | 4 | 8 | NA | NA | 768 | 1 | 38 | 5 |
| pSugE(p) | 14 | 14 | 10 | 2 | 8 | 4 | 8 | NA | NA | 512 | 1 | 38 | 4 |
| **MBEC | Mean MBEC (µg/ml) (n=6) |  |  |  |  |  |  |  |  |  |  |  |  |
| pMS119EH | 64 | 32 | 64 | 13 | 16 | 128 | 32 | 3750 | 64 | 4096 | 128 | 1024 | 4 |
| pEmrE | 256 | 64 | 256 | 32 | 106 | 32 | 48 | 7500 | 512 | 16384 | 256 | 2048 | 8 |
| pQacE | 32 | 64 | 256 | 13 | 32 | 32 | 32 | 7500 | 256 | 16384 | 512 | 2048 | 32 |
| pQacEΔ1 | 32 | 256 | 64 | 13 | 21 | 128 | 32 | 3750 | 64 | 16384 | 213 | 1024 | 8 |
| pQacF | 128 | 128 | 64 | 26 | 26 | 128 | 32 | 7500 | 149 | 16384 | 128 | 2048 | 21 |
| pQacG | 256 | 256 | 128 | 32 | 64 | 128 | 48 | 3750 | 64 | 8129 | 128 | 2048 | 8 |
| pQacH | 256 | 128 | 256 | 32 | 128 | 128 | 48 | 7500 | 512 | 16384 | 128 | 2048 | 32 |
| pGdx | 1024 | 256 | 32 | 13 | 64 | 128 | 24 | 6250 | 256 | 2048 | 128 | 2048 | 4 |
| pSugE(p) | 256 | 256 | 32 | 13 | 64 | 512 | 24 | 6250 | 171 | 16384 | 128 | 2048 | 8 |

\*10<sup>-3</sup> dilutions of plasmid transformed cultures adjusted to OD<sub>600 nm</sub> 1.0 as starting inoculum
\*\* 24 hr biofilms on MBEC device pegged lid incubated followed by drug incubation for 24 hrs in LB broth
NA; data not available- drug concentration absorbance values exceeded detection thresholds
**Antimicrobial abbreviations:** cetylpyridinium chloride (CPC), cetrимide bromide (CET), benzalkonium chloride (BZK), didecyldimethylammonium bromide (DDAB), cetyldimethylethylammonium bromide (CDAB), cetyltrimethylammonium bromide (CTAB), domiphen bromide (DOM), ethidium bromide (ET), methylviologen dichloride (MV), chlorhexidine dichloride (CHX), acriflavine (ACR), erythromycin (ERY), and tobramycin (TOB)

**Table 3.** Summary of AST MIC and MBEC values determined for *E. coli* KAM32 transformed with plasmids expressing various SMR genes grown as planktonic (broth), colony (agar spot), and biofilm (MBEC).

| Plasmid transformed | Mean MIC (µg/ml) n=3 |  |  |  |  |  |  |  |  |  |  |  |  |
| --- | --- | --- | --- | --- | --- | --- | --- | --- | --- | --- | --- | --- | --- |
| *Agar | CTP | CET | BZ | DDAB | CDEB | CTAB | DOM | ET | AC | MV | CHX | ERY | TOB |
| pMS119EH | 2 | 1 | 1 | 0.5 | 2 | 4 | 2 | 12 | 1 | 80 | 4 | 2 | 4 |
| pEmrE | **8** | **4** | 2 | **2** | **8** | **8** | **4** | **256** | **8** | 160 | 4 | 2 | 2 |
| pQacE | **8** | **8** | **4** | 1 | **8** | **8** | **4** | **32** | **4** | 80 | 4 | 2 | 2 |
| pQacEΔ1 | 2 | **4** | 1 | 0.5 | 4 | 4 | 2 | 8 | **4** | 80 | 4 | 2 | 4 |
| pQacF | **8** | **8** | **4** | **2** | **8** | **8** | **4** | 8 | **4** | 80 | 4 | 2 | 8 |
| pQacG | **8** | **8** | **4** | **2** | **8** | 4 | 2 | 8 | 1 | 80 | 4 | 2 | 8 |
| pQacH | **8** | **4** | **4** | **2** | **8** | **8** | **8** | **32** | **4** | 80 | 4 | 2 | 4 |
| pGdx | **8** | 2 | 1 | 0.5 | 4 | **8** | 2 | 16 | **4** | **320** | 4 | 2 | 4 |
| pSugE(p) | **8** | 2 | **2** | 0.5 | **8** | 4 | 2 | 16 | **4** | 80 | 4 | 2 | 4 |
| *Broth | Mean MIC (µg/ml) n=3 |  |  |  |  |  |  |  |  |  |  |  |  |
| pMS119EH | 1.2 | 2 | 2 | 0.5 | 2 | 1 | 1 | 16 | 8 | 80 | 1 | 1.5 | 2 |
| pEmrE | 2.4 | 4 | 2 | 1 | 2 | 1 | 1.5 | **64** | **64** | **320** | 1 | 1 | 2.5 |
| pQacE | 1.6 | 4 | 2 | 1 | 2 | 1 | 1.5 | **64** | 24 | 80 | 1 | 1.5 | 2 |
| pQacEΔ1 | 1.6 | 2 | 2 | 0.5 | 2 | 1 | 1 | 32 | 8 | 80 | 1 | 1 | 2 |
| pQacF | 1.6 | **8** | **4** | **2** | 2 | **2** | 1.5 | 16 | 8 | 80 | 1 | 2 | 2.5 |
| pQacG | 2.4 | 4 | **4** | **2** | 4 | **2** | 1.5 | 16 | 8 | 160 | 1 | 2 | 3 |
| pQacH | 1.6 | 4 | **4** | **2** | 4 | 1 | 1.5 | 32 | 24 | 160 | 1 | 1.5 | 2.5 |
| pGdx | 1.6 | 4 | 2 | 1 | 2 | 1 | 1.5 | 16 | 8 | 160 | 1 | 2 | 2 |
| pSugE(p) | 2.4 | 4 | 1 | 1 | 2 | 1 | 1.5 | 16 | 8 | 80 | 1 | 1 | 2 |
| **MBEC | Mean MBEC (µg/ml) n=6 |  |  |  |  |  |  |  |  |  |  |  |  |
| pMS119EH | 128 | 16 | 32 | 16 | 16 | 8 | 8 | 64 | 32 | 4096 | 32 | 64 | 3 |
| pEmrE | **32** | 32 | 16 | 16 | **64** | 8 | 26 | **512** | **256** | **16384** | 64 | 128 | 10 |
| pQacE | **32** | 32 | 16 | 8 | 53 | 8 | 16 | **256** | **213** | **16384** | 64 | 128 | 8 |
| pQacEΔ1 | 64 | **64** | 32 | 6 | 16 | 4 | 8 | 64 | 64 | **16384** | 64 | 85 | 8 |
| pQacF | 64 | **64** | 32 | 16 | 48 | 4 | **32** | **256** | 64 | 4096 | 32 | 64 | **16** |
| pQacG | 256 | **128** | 32 | 16 | 37 | 16 | 8 | 53 | 21 | **16384** | 64 | 64 | 8 |
| pQacH | **32** | 16 | 16 | 16 | 45 | 16 | **3** | **341** | **256** | 4096 | 64 | 128 | **27** |
| pGdx | **32** | **<8** | 32 | **4** | 13 | **32** | 13 | **341** | 64 | **16384** | 64 | 85 | 8 |
| pSugE(p) | **32** | 32 | **8** | 8 | 11 | 8 | 26 | **512** | 53 | **16384** | 64 | 64 | 4 |

\*10<sup>-3</sup> dilutions of transformant cultures adjusted to OD<sub>600 nm</sub> 1.0 as starting inoculum
\*\* 24 hr biofilms grown on MBEC device pegged lid incubated with drug for 24 hr in LB broth
**Antimicrobial abbreviations:** cetylpyridinium chloride (CPC), cetrime bromide (CET), benzalkonium chloride (BZK), didecyltrimethylammonium bromide (DDAB), cetyltrimethylethylammonium bromide (CDAB), cetyltrimethylammonium bromide (CTAB), domiphen bromide (DOM), ethidium bromide (ET), methylviologen dichloride (MV), chlorhexidine dichloride (CHX), acriflavine (ACR), erythromycin (ERY), and tobramycin (TOB)

Nearly all BW25113 SMR transformants examined using broth or agar AST methods had mean MIC values that were statistically insignificant (within a 2-fold change or identical) to the parental vector control BW25113-pMS119EH, with the exception of the pEmrE transformant (Table 2). BW25113-pEmrE transformants agar AST results showed increased QAC tolerance (MIC values ≥4-fold) to ethidium bromide (ET) and acriflavin (AC) as compared to vector controls (Table 2), in agreement with previous agar spot plated AST findings (54). All BW25113 SMR gene transformants grown as biofilms using the MBEC AST method demonstrated a significant increase in tolerance (≥4-fold change) to at least one QAC, inter-chelating dye, and/or antibiotic for each SMR transformant as compared to the parental vector control (Table 2). As expected from a previous study (28), BW25113 pQacEΔ1 transformant biofilms did not show significant MBEC values most antimicrobials we tested, with the exception of 4-fold higher MBEC values for cetrimide (CET) and methyl viologen (MV) as compared to the parental vector (Table 2). Interestingly, this less active *qacEΔ1* still confers some antimicrobial selection to specific QAC substrates in ‘wildtype’ BW25113 biofilms. BW25113 pEmrE transformant biofilms demonstrated the highest MBEC values to the broadest range of antimicrobials we tested as compared to vector controls and other pQac vectors tested (Table 2), once again supporting its designation as an ‘archetypical’ SMP member.

Lastly, BW25113 pGdx and pSugE(p) biofilms both conferred significantly higher MBEC values (≥4-fold change from vector control) to the fewest antimicrobials as compared to pQacE, pQacF, pQacG and pQacH transformants (Table 2), as expected for GDM and SMP subclass members (2, 4) confirming our first hypothesis. Similar to previous *E.coli* Gdx studies (11, 55), significant (4-fold) increased tolerance to CET and CTP was observed for BW25113 pGdx transformants but only during biofilm AST (Table 2). BW25113 pGdx transformant biofilm AST also demonstrated 4-fold increased MBEC values to previously unidentified QACs CDEB, and dye AC when compared to the parental vector control indicating biofilm AST may identify more Gdx-selective substrates (Table 2). BW25113 pSugE(p) biofilms also demonstrated 4-fold increased MBEC values for CTAB, MV, and TOB when compared to the ‘archetypical’ pGdx transformants and parental vector control, highlighting GDM substrate selectivity differences between closely related *sugE(p)* and *gdx* (Table 2).

Therefore, QAC tolerant phenotypes associated with SMR gene overexpression in a ‘wildtype’ *E. coli* strain background were most significant as biofilms. BW25113 biofilm AST also demonstrated broader antimicrobial tolerance profiles for each *qac* as compared to narrow substrate selection profile for pSugE(p) and pGdx transformants supporting hypotheses 1-2.

### 2.3 AST of E. coli SMR gene transformants show significant QAC tolerance as colonies and some QAC susceptible phenotypes as biofilms in an E. coli efflux pump deletion strain

A previous study comparing efflux pump gene over-expression in *E.coli* strains using agar spot plated AST methods, identified that mutations to the dominant Resistance-Nodulation-Division (RND) AcrAB efflux pump system were necessary to accurately measure MIC value of other secondary efflux pump systems, including EmrE (54). Therefore, we repeated AST experiments using *E. coli* strain KAM32, a common *E. coli* strain for efflux pump characterization, which lacks efflux pumps *acrB* and *mdtK* (Table 3; Fig. 1F). Due to the reduced growth and fitness of KAM32 we observed and as previously observed by (56), all SMR transformants resulted in lower mean MIC and MBEC values when compared to the BW25113 strain to all of the antimicrobial compounds we tested (Table 3), indicating that the loss of intrinsic efflux pumps enhanced antimicrobial susceptibility of KAM32. Agar spot plate AST of KAM32 transformed with pEmrE and all pQac vectors (except pQacEΔ1) demonstrated a significant increase (≥4-fold) in MIC values for most antimicrobials we tested when compared to the parental vector (Table 3; Fig. 1F). As observed for BW25113 SMR transformants, KAM32 transformants examined by broth AST demonstrated the least significant MIC values to all antimicrobials tested as compared to agar colony AST methods and highlights the inferiority of broth AST methods for QAC susceptibility testing (Table 3).

To ensure that a lack of SMR protein accumulation was not the cause of insignificant broth AST MIC values, we isolated the membranes (inner and outer membranes) from all BW25113 and KAM32 transformant broth cultures for sodium dodecyl sulfate (SDS)-Tricine polyacrylamide gel electrophoresis (PAGE) analysis (File S3A-B). SDS-Tricine (16%T) PAGE analysis confirmed that 12 kiloDalton bands corresponding to the estimated size of each over-accumulated SMR protein were detectable in both BW25113 and KAM32 transformant membranes as compared to membranes isolated from the pMS119EH transformant at the same IPTG induction conditions (Fig. S3). Furthermore, the possibility that efflux pump toxicity caused by protein over-accumulation within *E.coli* membranes due to IPTG induction was also not observed in BW25113 or KAM32 transformants based on OD_600 nm_ LB-AMP growth curves (Fig. S4). Therefore, AST growth are an important consideration for QAC tolerance determination, and broth microdilution AST methods appear to be least effective as compared to agar spot plating techniques to determine MIC values for SMR efflux pump overexpression in a KAM32 strain.

As compared to BW25113 transformant biofilms exposed to the same antimicrobials, KAM32 transformant biofilms resulted in fewer significant increased MBEC values as compared to parental vector controls (Table 3). Interestingly, KAM32 transformant biofilm AST MBEC results identified some SMR genes (*qacE*Δ*1, qacF, sugE(c), sugE(p)*) exposed to QACs (CET, BZK, DDAB, CTAB) had enhanced antimicrobial susceptibility (≤4-fold reduction in MBEC values) as compared to the vector control (Table 3). This indicates that when dominant efflux pump *acrB* and *mdtK* pump are absent in *E.coli*, the overexpression of SMR efflux pumps work against the cells making them more susceptible to the aforementioned QACs. To determine if SMR overexpression was toxic for biofilm growth, thereby reducing overall biofilm formation in BW25113 and KAM32 strains prior to antimicrobial exposure, the biofilm biomass accumulation on MBEC device pegged lids after 24 hr biofilm growth was quantified using CV staining (Fig. 2A-B). Both BW25113 and KAM32 transformants showed no significant reductions in CV stained biomass on the MBED device pegs when compared to their respective parental vector controls (Fig. 2A-B) indicating that SMR vectors did not significantly reduce biofilm formation. Interestingly, some KAM32 transformants had significantly increased biomass formation (1.5- to 2-fold) for pQacE, pQacEΔ1, pQacG, pQacH, pSugE(p) transformants as compared to pMS119EH (Fig. 2B) suggesting these transformants may promote biofilm formation. Hence, SMR transformants did not significantly impair biofilm biomass formation in either strain. To verify if the deletion of the dominant RND efflux pump *acrB* was an influential factor in biofilm biomass formation, BW25113 biofilm formation was compared to JW0451 (Δ*acrB*) and KAM32 was compared to its parental strain TG1 (Fig. 2C); no significant differences in biomass were observed for any strain pair in this study.

**Figure 2.**
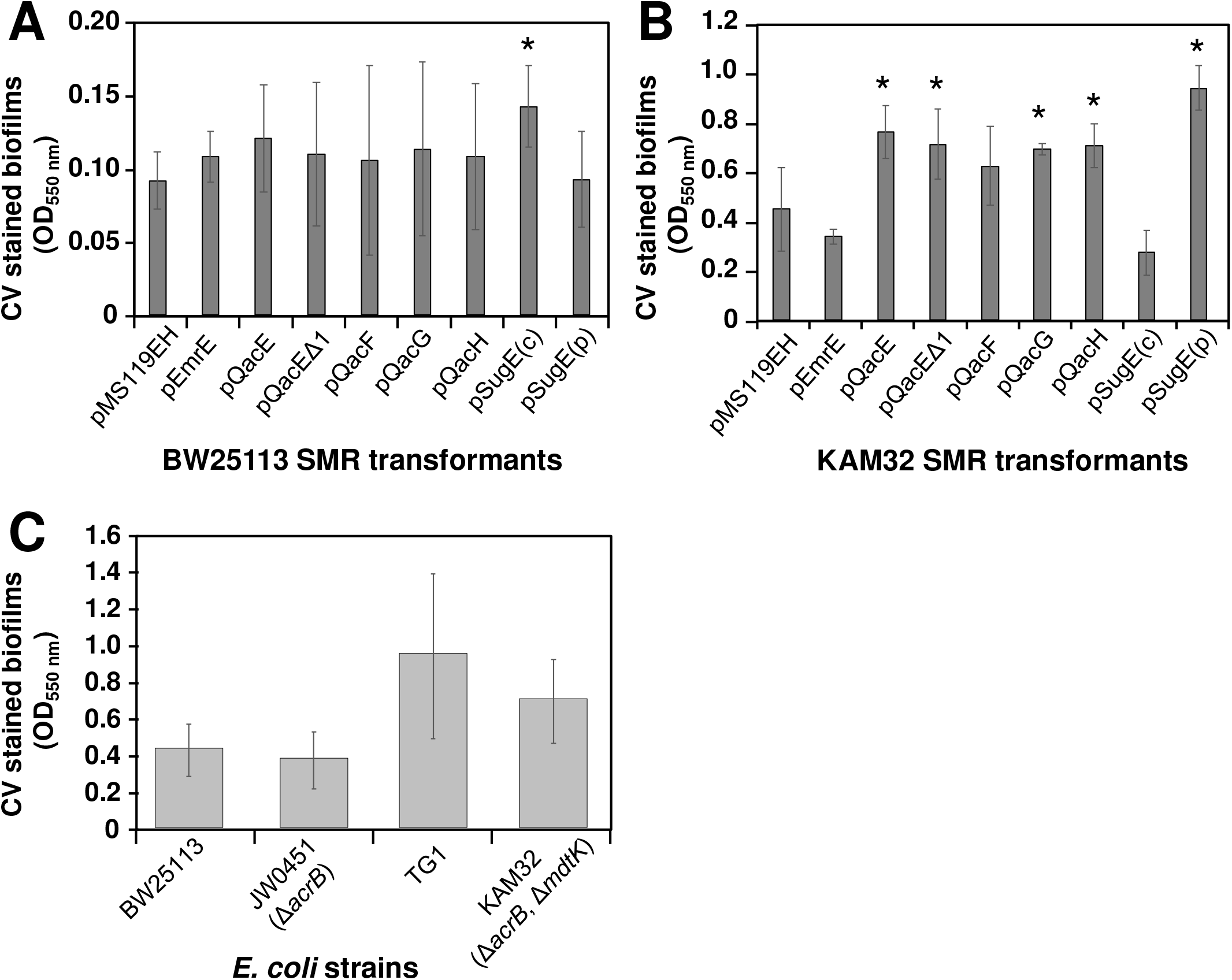
Biofilm biomass determination based on A_750nm_ CV stained values of MBEC device pegs of various *E. coli* strains and transformants. **A)** BW25113 SMR plasmid transformant biomass after 24 hrs at 37°C in LB medium. No significant differences in *p*-values ≤0.05 we determined for any samples as compared to the pMS119EH (parental vector) control strain. **B)** KAM32 SMR plasmid transformant biomass after 24 hrs at 37°C in LB medium. Asterisks (*) indicate significantly different p-values ≤0.05 as compared to pMS119EH (parental vector) control. **C)** Biofilm biomass (A750 nm) determination of wildtype strains *E. coli* BW25113, JW0451 (Δ*acrB*), TG1, and KAM32 (Δ*acrB*, Δ*mdtK*) strains grown for 24 hrs in LB medium at 37°C using MBEC devices. Significantly different p-values ≤ 0.05 (*) are noted by asterisks based on two-tailed Students T-tests.

Altogether, SMR gene expression in KAM32 increased the detection of significant MIC values by each SMR transformant when grown as agar colonies. The antimicrobial tolerance of various SMR transformants differed dramatically in KAM32 as compared to the wildtype BW25113, particularly when grown as a biofilm emphasizing the importance of growth conditions, and efflux pump deletions when performing AST. Comparing different *E.coli* strains also reveals that KAM32 can determine the broadest SMR substrates from AST profiles; however, caution should be taken when comparing agar colony versus biofilm values, since SMR expression in strains lacking *acrB* and *mdtK* increased susceptibility to some QACs when grown as biofilms.

### 2.4 Plasmid encoded SMR genes are regulated by either Type II Guanidinium riboswitches or class I integron promoters based on phylogeny

To determine how each SMR gene is regulated on their respective plasmids (and integrons), we examined the 500 nucleotide (nt) upstream region of each SMR gene starting from the +1 start codon (Tables S2-S3). The results demonstrated that all *qacE*Δ*1* and *qacE* genes primarily possessed a ‘Pq’ promoter sequence (identical to the sequence described by (15, 37)), −100 nt immediately upstream of +1 start codon (Fig. S5A,C). Remaining *qac* genes, *qacF, qacH*, and *qacG*, had very low (≤22%) or no sequence identity to either class I integron Pq within the 500 nt upstream region (Fig. S5B-C); the only exception was for 2 of the 91 *qacH* plasmids we examined had ≥90% sequence identity to Pq promoter (Fig. S5B). Expanding the inspection of nearby open reading frames on these plasmid maps encoding each *qac* gene showed that the majority (81-94%) of *qacF, qacH*, and *qacG* sequences were located within the variable cassette region of class I integrons or nearby transposable elements (Fig. S6). Hence, *qacF, qacH*, and *qacG* genes are most likely regulated by the class I integron *intI1* Pc promoter or to a lesser extent other promoter(s) associated with upstream transposable elements (Fig. S6). It is important to note, that the majority of Pc promoter sequences (56-94%) we identified from nearly all ‘qac’ containing plasmids surveyed had ≥90% sequence identity to the so-called ‘weak’ version of the class I integron promoter (Table S2; Fig. S8), similar to trends reported in a previous study (38). The remaining Pc sequences were hybrids of the string and weak Pc promoters, as described by Guerin *et al*. 2011 (38), or were not identified at all within the plasmids. Finally, a small proportion of all *qacE*Δ*1* gene encoding plasmids we surveyed (664 plasmids) had one or more additional SMR sequences located elsewhere on the plasmid, specifically, *qacF* (3/16; 18.8%), *qacG* (3/17; 17.6%), *qacH* (6/91; 6.6%) or sugE(p) (21/115; 18.3%) (Fig. S6). As a result, the expression of additional SMR genes are likely regulated by different plasmidic/integron promoters or riboswitch elements within the SMR gene.

Surveys of the 500 nt region upstream of *sugE(p)* encoding plasmids demonstrated high (≥85%) sequence identity to the *E. coli gdx* 5’ UTR of type II Gdm^+^ riboswitch for every *sugE(p)* regions we examined (Fig. S7A-B). This strongly suggests that *sugE(p)* gene are controlled by Gdm^+^ riboswitches supporting the third study hypothesis. Examination of other upstream regions for sugE sequences from Rhizobiales and Candidatus *Competibacter denitrificans* plasmids demonstrated poor riboswitch sequence identity to other known Gdm^+^ riboswitch types (Type I; *Clostridiales* Gdx WP_021653285.1, Type III; *Micromonospora* Gdx WP_013284696.1) and other proteobacterial species using BLASTn searches (Fig. S7B; Fig. S8). These α-proteobacterial *sugE* sequences carried on plasmids may be regulated by plasmid promoters associated with upstream genes or possess novel riboswitch regions.

### 2.5 Pq promoters associated with SMR genes show higher expression in planktonic versus biofilm cultures

Due to the high proportion of *qacE* and *qacED1* associations with the class I integron P*q* promoter and *qacF, qacG, qacH* regulation by Pc ‘weak’ promoters, we synthesized and cloned the P*q* (pPq) and Pc (pPc) ‘weak’ promoter regions (Fig. S8) into a *lacZ* reporter vector to characterize and compare promoter activity in *E. coli* BW25113 using β-galactosidase 96 well assays (Table 1). β-galactosidase activity of pPq and pPc was compared to control pMS119EH vector with (placZ; P*tac* promoter positive control) and without *lacZ* gene (pMS119EH; negative control) and induced with and without 0.05 mM IPTG (+/-IPTG) to match AST experiments. β-galactosidase activity of planktonic (broth) transformant cultures grown in rich medium (LB-AMP) sampled over the course of 24 hrs showed high β-galactosidase activity for both pPq and pPc transformants as compared to placZ+IPTG controls up to and including 11 hrs (Fig. 3A). At 24 hrs, all planktonic transformants (excluding pMS119EH) demonstrated reduced levels of β-galactosidase activity similar to placZ-IPTG transformants, which was anticipated as cultures reach stationary phase. At early to mid-log growth phases (3-7 hr timepoints), planktonic pPc transformants had reduced (50-75%) β-galactosidase activity as compared to pPq, suggesting Pq is a much stronger promoter than Pc under these conditions. In fact, pPq transformant β-galactosidase activity was similar to placZ+IPTG control values in most analyses (Fig. 3C-E) suggesting that pPq expression is similar to the 0.05 mM IPTG induced Ptac expression levels used for all AST experiments in this study. In comparison, *E.coli* transformants grown as biofilms showed significant reductions in β-galactosidase activity for pPq (35.3%) and pPc (41.1%) as compared to placZ+IPTG (Fig. 3B) and indicate that Pq and Pc promoter activity is lower in an established 24 hr biofilm. Comparing β-galactosidase activity from pPq and pPc transformant biofilms to values obtained for stationary (24 hr) planktonic pPq and pPc transformants, we observed a similar Miller unit expression levels. This suggesting that 24 hr biofilms obtained from pegs had similar vector expression levels as stationary phase planktonic *E. coli* (Fig. 3A-B).

**Figure 3.**
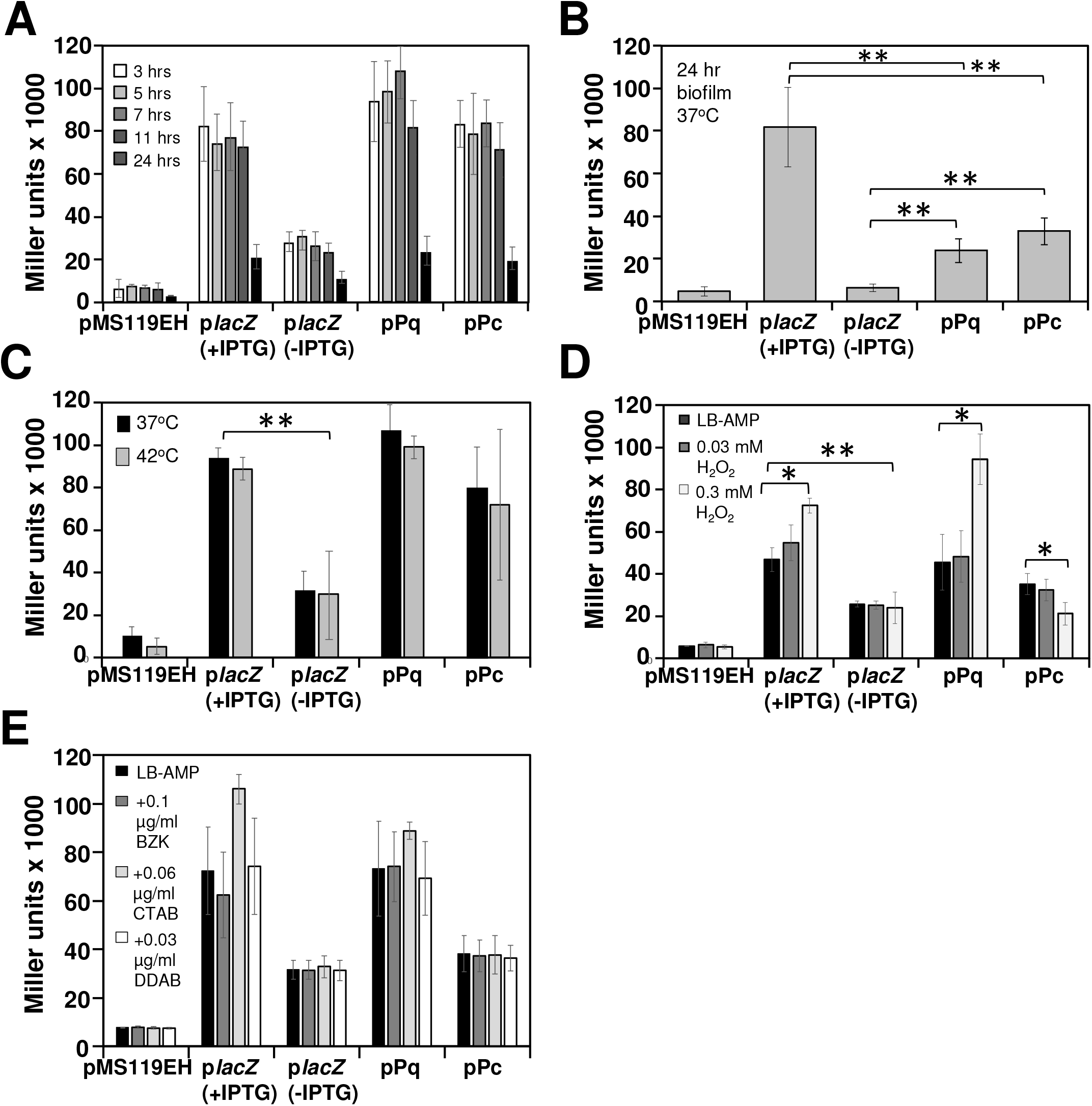
β-galactosidase assays of *E. coli* BW25113 transformed *lacZ* reporter in LB medium to determine promoter expression differences as planktonic and biofilm cultures. Refer to legends in panels for transformants and exposure conditions tested. **A)** Miller units calculated for BW25113 transformants assayed at 3, 7, 11 and 24 hrs of growth. **B)** Miller units determined for BW25113 transformants grown as biofilms after 24 hrs of growth at 37°C. Miller units for BW25113 transformants grown planktonically in LB broth exposed to **C)** 37°C and 42°C for 5 hrs, **D)** 0, 0.03, and 0.3 mM H_2_O_2_ for 5 hrs and **E)** sub-inhibitory concentrations of 0.1 μg/ml BZK, 0.06 μg/ml CTAB, 0.03 μg/ml DDAB for 5 hrs at 37°C. All samples shown in panels A-E were measured in triplicate and asterisks shown on bars indicate significant p-value differences of ≤ 0.01 (**) and ≤ 0.05 (*) according to two-tailed Students T-test comparisons. In all panels, a final concentration of 0.05 mM IPTG was added to placZ + IPTG samples.

### 2.6 Pq and Pc promoters are differentially regulated by various stress conditions in planktonic cultures

To verify what stress conditions induced Pq and Pc promoter expression, we examined planktonic transformants in greater detail. Mid-log phase (5hrs; OD_600nm_ = 0.4-0.7 units) transformant cultures grown in LB-AMP broth were exposed to various stressors: high temperature (42°C), H_2_O_2_-induced oxidative stress, and sub-inhibitory QAC exposure (CTAB, DDAB, and BZK) (Fig. 4C-E). β-galactosidase activity determined for both the pPq and pPc transformants showed small but statistically significant increases in expression in the presence of high concentrations of H_2_O_2_ (0.3 mM; Fig. 4D), indicating that both Pq and Pc promoters are regulated by oxidative stress. QACs are predicted to enhance oxidative stress as part of their mechanism of action (6) and unfortunately, we did not observe and significant differences response to sub-inhibitory QACs by pPq and pPc transformants in LB (Fig. 4D-E). This result may be due to the complexity of the rich LB medium which may absorb QACs at sub-MIC concentrations; therefore, we repeated β-galactosidase assay in minimal medium (M9-AMP). Switching to M9 growth increased β-galactosidase activity differences between IPTG-induced and uninduced transformant control vectors made this minimal media more ideal for distinguishing pPq and pPc expression differences (Fig. 4A-E). Similar to LB-AMP, sampling β-galactosidase activity in M9-AMP over 24 hrs showed higher activity in both pPq and pPc sampled at 3-11 hrs (Fig. 5A). Only pPc transformants in M9-AMP demonstrated a small but statistically significant increase in 42°C heat induced expression of the Pc promoter (Fig. 5B) suggesting heat stress may induce the ‘weak’ Pc promoter. Both pPq and pPc transformants showed statistically significant increases in expression when exposed to 0.03 mM H_2_O_2_ as compared to no H_2_O_2_ exposure. Unfortunately, exposure to 0.3 mM was toxic to all M9-AMP cultures so data could not be collected (Fig. 5C). Only pPq transformants in M9-AMP demonstrated a small (10-14%) but significant increase β-galactosidase activity when exposed to QACs (CTAB and DDAB; Fig. 5D-F), suggesting that the Pq promoter is more sensitive to QAC-induced damage by comparison to the Pc promoter grown planktonically. Therefore, Pq and Pc promoters respond to various oxidative stressors during planktonic growth conditions, where the Pc promoter is more responsive to H_2_O_2_-induced stress and to lesser extent heat-induced stress.

**Figure 4.**
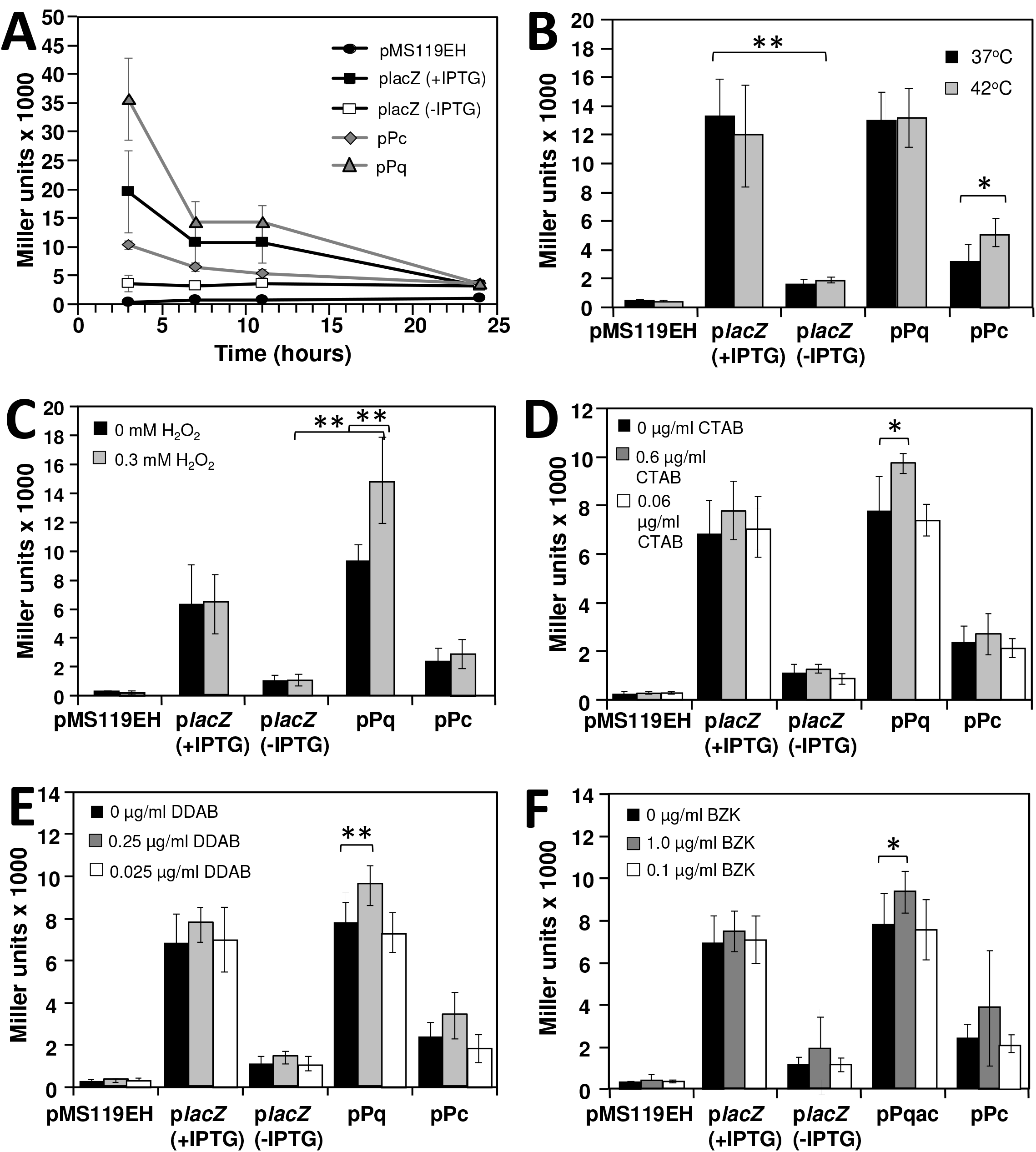
β-galactosidase detection assays of *E. coli* BW25113 *lacZ* reporter transformants grown in minimal M9 medium broth to determine promoter expression differences in planktonic cultures. Refer to legends in panels for transformants and exposure conditions tested. **A)** Miller units calculated for BW25113 transformants assayed at 3, 7, 11 and 24 hrs of growth. Miller units of BW25113 transformants exposed to **B)** 37°C or 42°C for 5 hrs, **C)** 0 or 0.3 mM H_2_O_2_ for 5 hrs, **D)** sub-inhibitory concentrations of CTAB for 5 hrs at 37°C and **E)** sub-inhibitory concentrations of DDAB (refer to panel legend) for 5 hrs at 37°C. All transformants were measured in triplicate and asterisks shown in bars on panels B-E indicate significant p-value differences of <0.01 (**) and <0.05 (*) according to two-tailed Students T-test comparisons. In all panels, a final concentration of 0.05 mM IPTG was added to placZ + IPTG samples.

## 3. Discussion

### 3.1 SugE/ Gdx sequence diversity is greater than Qac members and validates antimicrobial substrate selection differences between SMR subclasses

Two major findings were apparent from the bioinformatic survey and phylogenetic analysis of SUG protein sequences encoded on Gram-negative bacterial plasmids: 1) GDM members appear to distribute among more diverse phyla (Fig. 1A-C) and 2) GDM sequences have much greater sequence variability than Qac members (Fig. 1D, E; Fig. S2). High sequence variation among SugE members (Fig. 1E; Fig. S2) may suggest *sugE* horizontal gene transfer to plasmids originate from different bacterial phyla/species or are subject to lower selection pressure to maintain *sugE* sequences but in Enterobacteriales SugE(p) sequences appear to predominate over others (Fig. 1B-C). We speculate that GDM gene transfer between α-proteobacterial plasmids may biased in favor of the species they co-habitat with, since SUG sequences we identified on α-proteobacterial plasmids failed to identify *gdx/ sugE* homologs at significant BLASTn e-values (≤1.0 x10^−3^) in the respective genomes the plasmid was isolated from. Why GDM sequence diversity on plasmids is so different remains is unclear. GDM gene transfer and mobility on plasmids may be driven by selection pressure for Gdm^+^ containing substrates, as many Gdm^+^ containing chemicals like QACs are used in high quantities (100 kilotons annually) in food, agricultural, petrochemical, and clinical industries (57–59). GDM homologs have also demonstrated involvement in tolerance to metal containing antimicrobials such as di- and tri-butyltin based on previous studies of *Aeromonas molluscorum sugE* (60), suggesting some GDM members may recognize more than just the nitrogen cation similar to the ability of EmrE to recognize compounds with phosphorous centers (eg. tetraphenylphosphonium (as reviewed by (4)). Riboswitch type and activity in species that inherit the plasmid may also be a factor driving *sugE/ gdx* gene mobility. Altogether, the close sequence association between *qac* genes (particularly *qacE*Δ*1*) and *sugE(p)* make these SMR sequences useful genetic markers to monitor as proxies for QAC tolerant phenotypes and QAC pollution monitoring.

In contrast to plasmid encoded GDM sequences, Qac SMP homologs showed very high sequence identity to each other (Fig. 1D), but much lower identity to *E. coli emrE* (Fig. 1A; Fig S2), suggesting a single origin for plasmid *qac* sequences surveyed thus far. It remains unclear why *qacEΔ1* predominates over all other SMR sequences, including its original and functional progenitor *qacE* sequence (Fig. 1B-C). We speculate that the predominance of the less active QacEΔ1 may be more beneficial to overall cell fitness, when compared to fewer ‘functional’ Qac sequences we observed in the surveys i.e. QacF, QacG, and QacH, Evidence for this may be seen in our planktonic *E.coli* SMR transformant over-expression data (Fig. S4) and from KAM32 Qac transformant biofilm AST results (Table 3). Additional QAC mechanisms of tolerance may be larger influencing factor, such as the involvement of other intrinsic and unrelated QAC selective efflux pumps like MdtM (61) or bacterial adaptation to QACs resulting in porin, peptidoglycan and lipid biosynthesis alterations as noted in QAC adaptation studies of *E.coli* (62–64).

### 3.2 AST growth conditions and other efflux pump losses impact QAC tolerance conferred by SMR members

Overall, AST of 6 representative plasmid transmitted SMR transformants in *E.coli* K-12 strains resulted in moderate (4-fold increase from parental vector control) increases in MIC values for various QACs (Tables 2–3). *E.coli* ‘*qac*’ transformants (excluding pQacED1) grown as biofilms recognized the broadest amount of QACs as compared to Gdx and SugE, supporting/reconfirming the hypothesis that SMR subclass affiliation predicts SMR subclass affiliation. Furthermore, our QAC AST analyses support SMR substrate selection profiles reported in previous AST studies of QacE (28, 29, 31), QacEΔ1 (28, 29, 31), and QacF (30, 65), and Gdx/ SugE (10, 11, 66) indicating some reproducibility to SMR phenotypes. Our AST experiments also suggest that the mode of bacterial growth significantly impacts MIC values (Tables 2–3). Based on our AST findings, future QAC substrate selectivity testing of SMR genes and other QAC selective efflux pumps should avoid making substrate section determinations for planktonic AST methods or include additional colony/ biofilm growth AST methods. Our findings are in accordance when considering QAC mechanisms of action and preferred usage to eradicate biofilms; QACs are known to damage membranes and increase oxidative damage within cells (6, 67). Due to the multiple mechanism of action and bacterial killing at low QAC concentrations, QACs are frequently used by many industries to eradicate biofilms (33, 68), but low level QAC accumulation may inadvertently increase selection pressure to spread and express SMRs and other QAC-selective efflux pumps, particularly for improving biofilm growth. Therefore, planktonic growth to determine QAC tolerance by AST may be a misleading technique for estimating bacterial QAC tolerance and movement towards biofilm AST methods may better reflect bacterial QAC tolerance under physiologically relevant growth conditions.

The AST results we obtained for broth microdilution and agar dilution methods to determine MIC values for pQacE, pQacEΔ1, and pGdx members were consistent with previous studies (29, 31) that reported low-moderate increases in MIC values (2-4 fold MIC increases from wildtype) for QACs and inter-chelating dyes we included in our study. We have not only reconfirmed QAC selectivity for various SMRs in our findings, but also identified new QACs as SMR substrates. Based on the lower fold change differences in MIC values from our AST analyses, we suggest that plasmid transmitted SMR contributions to overall QAC tolerance in *E. coli* (and likely other Enterobacterales species) is likely low-level at best and require the host species to have other intrinsic QAC tolerance mechanisms in place for higher level antiseptic tolerance (>4-fold MIC values). For example, *E.coli* possess a number of additional unrelated QAC selective efflux systems (as reviewed by (1)) that may be more effective when expressed together during QAC exposure or during different growth physiologies as suggested in a previous study (54). The results of this study also provide the first phenotypic AST characterization of SMR genes *qacF, qacG*, and *qacH* which recognize and confer tolerance more QACs than QacE (Fig.1F). We show that pQacEΔ1 transformants confer tolerance to CET and AC (Table 2–3), despite the disruption of the 4^th^ transmembrane helix as noted in a previous study (28). These finding are helpful for ongoing surveillance studies and pertinent for *in silico* predictions of QAC tolerance when monitoring specific *qac* members. Hence, SMR genes remain a useful proxy for estimating QAC anthropogenic pollution and low-level antiseptic tolerance (13, 69).

Finally, AST results we obtained for KAM32 ‘Qac’ transformants demonstrated the most significant increase in MIC values to broadest range of QAC structures when grown as planktonic and agar colonies (Table 3) but fewer QACs were consistently identified as having a significantly enhanced or reduced MIC or MBEC values for all three AST growth conditions when compared (Fig. 1F). We suggest two factors that may have influenced these discordant KAM32 AST results between growth methods. 1) Different antimicrobial diffusion rates in liquid versus solid media may impact the antimicrobial agent concentrations exposed to cells during different grown states. Solid agar medium plates have lower antimicrobial diffusion rates as compared to liquid planktonic (high antimicrobial diffusion) and biofilm cultures. Biofilms are expected to have slower antimicrobial diffusion rates as compared to planktonic cultures as cells since biofilms are surface attached cells on pegs with more protection from the solid surface they attach to as reviewed by (33, 34). This might explain why agar plating KAM32 transformants identified significantly increased MIC values for more SMR transformants and antimicrobial combinations we tested when compared to MBEC devices or broth microdilution AST methods. Since media compositions for planktonic and biofilm AST were identical, our results suggest the mode of growth mode plays an influential role as well as the genetic background of the strains we tested (KAM32 versus BW25113).

The second factor influencing KAM32 AST result discordance, may pertain to SMR proteins’ ability to function as a ‘dual-topology’ dimeric transporter in the plasma membranes of a bacterium (as reviewed by (48, 70)). A dual-topology dimer refers to the ability of a transporter to fold and insert in either insertion orientation in a membrane, where the amine- and carboxyl-termini of the SMR protein can orient to face cytoplasm or the periplasm but form an asymmetric functional dimer. We observed that some KAM32 SMR transformant biofilms exposed to CTP and CET exhibited increased susceptibility (4-fold reduced MIC values) as compared to the vector control (Table 3), suggesting some SMR efflux pumps acted as importers for specific antimicrobials rather than exporters of these drugs. Due to the dual-topology nature of SMR protein dimers, previous studies of *E. coli* EmrE and SugE/ Gdx have shown an ability to act as antimicrobial importers depending on amino acid sequence alterations and protein folding conditions (66, 71). It may be possible that the loss of both *acrB* and *mdtK* in KAM32 grown as a biofilm creates membrane conditions favorable for SMR to function as an ‘importer’. Further examination into SMR folding in *E.coli* strains lacking efflux pumps *acrB* and *mdtK* grown as biofilms is needed to clarify this interesting observation.

### 3.4 SMP expression is regulated by integron promoters while GDM members are regulated Type II Gdm^+^ riboswitches

A recent study by Kermani *et al*. 2018 (2), stated that most SMR transmitted by plasmids relied on plasmid promoters. Our findings indicate this statement only applies to ‘qac’ SMR subclass homologs, whereas *sugE(p)* and perhaps other GDM members are regulated by a riboswitch. The findings for our survey show Enterobacterales *sugE(p)* predominate over all other *gdx/ sugE* sequences, therefore, it was no surprise that the type II Gdm^+^ riboswitch was detected similar to *E.coli gdx* riboswitch (Fig. S7). Translational control of plasmid transmitted *sugE(p)* sequences by riboswitches may explain why gdx/sugE sequences have more sequence diversity and origins. The correct species-riboswitch combination may be an additional level of gene expression control on Enterobacteriales plasmid gene expression, particularly under circumstances where *sugE(p)* expression may be controlled by other transcriptional elements and plasmid promoters.

Additionally, our findings identified that only *qacEΔ1* and *qacE* frequently associate with class I integron Pq promoters, whereas *qacF, qacG, qacH* were most frequently regulated by the integrase *intI1* Pc ‘weak’ promoter region (Table S2, Fig. S5, S6). Our observations that the ‘weak’ version of the Pc promoter predominates over ‘strong’ and ‘hybrid’ Pc sequences match findings reported by Guerin *et al*. 2011 (38) and suggest the popularity of the ‘weak’ Pc version may help offset the fitness costs to the organism caused by ‘strong’ Pc promoter gene cassette over-expression. High-level over-expression of the *qac* genes we cloned downstream from the Ptac promoter in the expression vector pMS119EH caused growth reductions at final concentration greater than 0.05 mM IPTG (Fig. S4). Fitness costs caused by potentially toxic gene over-expression from integrons may also explain why both the Pq and Pc integron promoters had β-galactosidase expression levels that were similar or lower than the 0.05 mM IPTG Ptac induction levels we selected as a positive control for SMR gene over-expression (Fig. 3). We suggest that gradual toxic over-expression caused by ‘leaky’ Pq promoters may explain why *qacE*Δ*1* predominates over *qacE*, as *qacE*Δ*1* has some efflux activity. Finally, gene over-expression fitness costs may explain why other functionally active *qac* gene isoforms are less abundant and regulated by ‘weak’ Pc promoters.

Our characterization of both pPq and pPc promoters grown as either planktonic or biofilm cultures after 24 hrs in LB medium demonstrated similar β-galactosidase expression levels (Fig. 3A-B). Examination of Pq and Pc promoters in M9 demonstrated that Pq promoters had higher β-galactosidase activity than Pc promoters (2-4-fold Miller units increases) and Pq were inducible by peroxide and sub-inhibitory QAC exposure, whereas Pc promoters showed slightly higher β-galactosidase activity when exposed to 42°C heat stress (Fig. 4). The conditions that induced Pq promoter in our β-galactosidase assays make sense when considering known QAC mechanisms of action that generate oxidative damage to proteins and DNA (6). However, previous Pc promoter studies indicate the promoter is regulated by the SOS response system, where UV light and DNA damage are primary inducers of the response regulator LexA which regulates and binds to the Pc promoter (72). SOS regulation may explain why we observed only modest increases in Miller units for peroxide and QAC exposure of pPc transformants in M9 medium. The shorter length of the synthesized ‘weak’ version Pc promoter region we synthesized for experimental analysis in this study may another influencing factor. Previous Pc promoter studies typically examined DNA regions 2-4 times the size of our selected core Pc promoter region (37, 38, 73). We avoided using longer Pc regions for analysis due to lower sequence identity between DNA regions surrounding the conserved Pc region core in our plasmid sequence dataset. Future studies of Pq and Pc regions will ideally focus on examining more diverse promoter regions beyond the core Pc region.

In conclusion we have shown that SMR gene distribution, sequence variation, and antimicrobial susceptibility on plasmids match trends identified for chromosomally inherited SMR members as noted in recent studies (2, 5). Our analysis support the hypotheses that SMR phylogeny and amino acid sequence can predict antimicrobial susceptibility profiles and particularly when grown as a biofilm in strains with functional intrinsic efflux pumps. Our analysis provides AST characterization of the most frequently detected SMR sequences transmitted on MDR plasmids and reveals SMR gene regulation appears to match chromosomally inherited SMR gene regulation trends. Altogether, this information provides more context to ongoing antimicrobial resistance genetic surveillance studies by providing QAC phenotypes to formerly uncharacterized genes and identifying optimal growth conditions for AST. This study also provides useful insights into how AST growth methods influence QAC antimicrobial phenotypes related to SMR over-expression and highlight a possible role for AcrB and MdtK in regulating SMR pump activity. Future studies will ideally explore how SMR genes contribute to multidrug resistant phenotypes with intrinsic efflux pump systems and other resistance genes carried on MDR plasmids.

## 4. Materials and Methods

### 4.1 SMR sequence collection and multiple sequence alignments

Plasmid-borne SMR protein and nt sequence searches were conducted using NCBI (https://www.ncbi.nlm.nih.gov/), UniProt (https://www.uniprot.org/), INTEGRALL (74), and BacMet (75) databases according to sequence annotation as ‘qac’ (*qacE, qacEΔ1, qacF, qacG, qacH, qacI, qacJ, qacZ*), ‘*sugE*’ or ‘smr’ and based on tBLASTn sequence similarity searches (76) to EmrE (NP_415075), QacE (NP_044260) and SugE (NP_418572) query sequences. Sequences related to PSMR members *E. coli* MdtIJ (MdtI; NP_416116 and MdtJ; NP_416117) were not identified from tBLASTn searches with significant e-values (≤ 1.0 × 10^−4^) to any plasmids and as a result are not discussed further. A total of 1463 Gram-negative proteobacterial plasmids with identified SMR protein sequences were obtained, where only 533 plasmid sequences had uniquely different accession sequence numbers. SMR protein sequences were further aligned using the online server COBALT (77) (Table S1; Fig. S1), where it was determined that only 20 SMR protein sequences had unique sequence identity (< 99%; Fig. S2). Unique SMR proteins were annotated as QacE, QacEΔ1, QacF/I, QacG, QacH, and SugE using multiple sequence alignment software Jalview version (v) 2.10.5 (78)

The 500 nt upstream region of frequently identified SMR gene sequences were also examined (Tables S2-3). Plasmid sequence accession numbers were obtained from either UniProt or GenBank and aligned relative to each SMR gene +1 reading frame start site (ATG/GTG) using EMBL-EBI Clustal Omega software (ClustalO; https://www.ebi.ac.uk/Tools/msa/clustalo/) linked to the Jalview software package (78). ClustalO nt alignments were used to identify and compare sequence identity of Pq and Pc promoter regions as well as Types I-III Gdm^+^ riboswitch consensus sequence identity summarized in Fig. S8. If no identity/ consensus was determined within the 500 nt upstream SMR sequence (as was the case for *qacF, qacG*, and *QacH*), manual SMR sequence locations were determined to predict the promoter region on the plasmid. Plasmid sequences surveyed are listed in Table S2-S3 and include Pq and Pc promoter detection by the BLAST+ software package (79) to perform BLASTn searches of various query sequences (listed in the tables) and the outcomes are summarized on Tables S2-S3. Due to the excessively large plasmid alignment file sizes (>14 MB each), only plasmid sequence accession numbers are provided in Tables S2-S3.

### 4.2 Phylogenetic analysis of SMR proteins

Phylogenetic analysis of the 20 unique SMR protein sequences was performed using the online PhyML Maximum Likelihood estimation methods (PhyML; (80–82); http://www.atgc-montpellier.fr/phyml/), with 100 bootstrap replicates to determine branching confidence at specific nodes. During PhyML estimations, an approximate Likelihood Ratio test and approximate Bayes estimation methods were used, both resulting in similar clusters; aBayes analysis is shown in Fig. 1A. *E. coli* EmrE NP_415075 and SugE NP_418572 sequences represented archetypical SMP and SUG subclass members respectively, and *Archaeoglobus fulgidus* QacE sequence AAB89552 served as the outgroup for analyses (Fig. 1A).

### 4.3 Homology modelling

Homology models of QacE (NP_044260) and SugE(p) (YP_002302254) protein sequences were generated using the online server I-TASSER (83). QacE and SugE(p) sequences showed the most significant homology based on C-scores of 0.12 and 0.22 respectively to the *E. coli* EmrE X-ray crystal structure monomer A (3b5dA.pdb; (48)). I-TASSER homology models were used to generate the protein ribbon cartoon images in Fig 1D-E using Pymol v.1.3 software (84).

### 4.4 Drugs, strains, plasmids, and cloning procedures used in this study

Antimicrobial compounds tested in this study were obtained from Tokyo Chemical Industry Co. (Portland, OR, USA) or Millipore-Sigma (Burlington MA, USA) and prepared in sterile distilled water at 50 mg/ml stock concentrations (unless otherwise specified): cetylpyridinium chloride (CPC), cetrimide bromide (CET), benzalkonium chloride (BZK), didecyldimethylammonium bromide (DDAB), cetyldimethylethylammonium bromide (CDAB), cetyltrimethylammonium bromide (CTAB), domiphen bromide (DOM), ethidium bromide (ET), methylviologen dichloride (MV), 0.25 mg/ml chlorhexidine dichloride in 95% ethanol (CHX), acriflavine (ACR), 20 mg/ml erythromycin (ERY), and 20 mg/ml tobramycin (TOB).

Plasmid-encoded SMR sequences representing each annotated subclade in Fig. 1A-C (ie. *qacE, qacED1, qacF*, *qacG*, *qacH* and *sugE* (p)) as well as promoter region sequences, Pc and Pq, shown in Fig. 2A were selected for further experimental analysis involving gene synthesis and cloning into pUC-57 by BioBasic Inc. Gene Synthesis Services (Markham, Ontario, Canada; Table 1). SMR genes were individually subcloned into the multiple cloning region 5’ XbaI and 3’ HindIII (or 3’ PstI) restriction sites of the low-copy vector pMS119EH (50) for IPTG inducible Ptac promoter expression using cloning procedures described by (85). A detailed description of the cloning procedures to construct each *lacZ* reporter construct with upstream P*tac*, P*c*, and P*q* promoter regions is summarized in File S1 and involved a modified version of the pMS119EH vector. All vectors were constructed and manipulated in *E. coli* strain DH5α and all vectors were sequence verified using Sanger sequencing services from Eurofins Genomics (MWG Operon CA, USA). Forward and reverse sequencing primers for pMS119EH, *Ptac*, and Trp repressor regions are listed in File S1.

Previously cloned SMR genes from *E. coli emrE* (86) and *sugE* (11), were also included in this analysis in pMS119EH, as representative SMP and SUG family members for comparative analysis. Plasmids were transformed into *E. coli* K-12 wildtype BW25113 strain (51) as well as an efflux pump deficient strain, KAM32 (*acrAB*, *mdtK*; (52); Table 1), for all AST described below. SMR protein accumulation from extracted cell membranes of plasmid transformed BW25113 and KAM32 strains was performed and confirmed using 16% (T) sodium dodecyl sulfate (SDS)-Tricine PAGE analysis using the *E. coli* cell membrane extraction procedure described by (56); gel loading details are provided in Fig. S3.

### 4.5 Broth microdilution AST method

All AST analyses were performed using overnight (18 hr) cultures of each transformant grown in Luria Bertani (LB) broth with 100 μg/ml final concentration ampicillin (LB-AMP) were used to inoculate 96-well microplates for AST at 37°C with shaking at 155 rpm. Overnight cultures were standardized to an optical density at 600 nm (OD_600nm_) of 1.0 unit prior to their final dilutions for each method. Broth microdilution AST of plasmid transformed *E. coli* strains (BW25113 and KAM32) was performed as described previously (87). Briefly, standardized overnight cultures were diluted 10^−3^ into fresh LB-AMP medium containing log2 dilution gradients of 1 of 12 antimicrobial stock solutions (listed in section 4.4) in each row of a 96-well Nunc flat-bottomed polystyrene microplate. Optimal expression of *qac* and *gdx/sugE* genes from our inducible vector was determined from growth curve analysis after 24 hrs in the presence of 0.05 mM (final in-well concentration) of IPTG in LB-AMP broth (Fig. S4). Each microplate had wells containing uninoculated media and antimicrobial drug concentration as baseline controls. Growth was based on OD_600nm_ measurements using a Multiskan Spectrum Ultraviolet /Visible wavelength region microplate reader (ThermoScientific, Waltham, MA). Three biological replicates of each strain were tested in duplicate (n=6) and statistically assessed using two-way student’s T-test to determine significant *p*-value differences as compared to the parental vector. MIC values for each transformant were determined to be the lowest antimicrobial concentration that resulted in a lack of growth (OD_600nm_ value) that was indistinguishable from uninoculated media containing drug control well.

### 4.6 Agar (spot) dilution AST method

Agar dilution AST of SMR plasmid transformants (BW25113 or KAM32) was performed as described previously (87) on LB-AMP agar petri plates to maintain expression vectors. Overnight cultures were standardized as described for broth microdilution AST and were diluted to 10^−3^ for spot plating. A sterilized stainless steel 48-pinned replicator was used to spot 1 μl of diluted culture onto LB-AMP agar plates, each plate containing a log2 serial dilution of the antimicrobial stock concentration and 0.05 mM IPTG. Agar dilution plates were incubated overnight for 24 hrs and in some cases 48 hrs at 37°C. Growth of each spot was scored by visual inspection, where the mean MIC values for each transformant was determined to be the minimum drug concentration that resulted in no visible colony growth. Table 2–3. A minimum of three biological replicates were measured for each transformant and compared to the respective strain transformed with the parental pMS119EH vector (Tables 2–3).

### 4.7 Biofilm AST and analysis methods

Minimum biofilm eradication concentration (MBEC) determination experiments of SMR gene containing plasmid transformed BW25113 and KAM32 *E. coli* was performed as described by (88) with the following modifications. Overnight cultures were sub-cultured twice on LB-AMP agar plates at 37°C for 18 hrs. Isolated colonies were resuspended in LB broth to McFarland standard of 1.0. Standardized cultures were diluted 10^−3^ into 180 μl LB-AMP broth containing 0.05 mM IPTG final concentration in 96-well microplates with an inserted 96-pegged lid MBEC assay device (Innovotech, AB). MBEC device plates were incubated at 37°C with shaking at 155 rpm for 24 hrs. The 24 hr biofilm coated pegged lids were rinsed twice in sterile phosphate buffered saline (PBS) and either stained with 0.5% (w/v) crystal violet (CV) for biomass determination as described by (88) or aseptically transferred into a new 96 well microplate containing fresh sterile LB-AMP with log2 dilutions of antimicrobial stock solutions for MBEC determination. MBEC determination experiments were incubated at 37°C with shaking (155 rpm) for another 24 hrs where, any biofilms that remained after drug exposure were harvested by aseptically washing the lids in sterile PBS twice and transferred to a new 96-well plate containing 200 μl recovery medium (sterile LB with 0.01% v/v Tween-20, 5×10^−3^ g/ml L-histidine, 5×10^−3^ g/ml L-cysteine, 1×10^−2^ g/ml reduced glutathione) followed by sonication of the pegs for 10 minutes using a Branson 3800 sonicating water bath. Sonicated biofilms recovered from the pegs in the 96 well plate were spot plated onto LB-AMP agar plates using a Boekel Scientific™ 48-steel pin replicator (1 μl transfer volume/ pin) and incubated for 24 hrs at 37°C to determine cell viability/ peg based on colony formation; the lowest antimicrobial concentration that prohibited colony formation was defined as the MBEC value. A total of 3 biological replicates were measured for each plasmid transformed strain. A Students T-test (2 tailed, paired, heteroskedastic) calculation was used to determine statistically significantly different MBEC values for SMR plasmid transformants as compared to the parental vector (pMS119EH) strains.

### 4.8 β-galactosidase reporter assays of planktonic and biofilm cultures

Planktonic LB-AMP broth cultures of plasmid-transformed *E. coli* BW25113 with placZ, pPc and pPq (listed in Table 1) were grown at various temperatures (either 37°C or 42°C), to various time points (6-48 hrs) as indicated with or without sub-inhibitory concentrations of antimicrobial using the high-throughput 96-well microplate β-galactosidase assay method described by (89). The only modification in this procedure was the replacement of PopCulture^®^ (EMD Millipore) solution with Fastbreak™ cell lysis reagent (Promega, WI, USA) solution to permeabilize cells prior to the addition of the β-galactosidase reaction-buffer (60 mM Na2HPO4, 40 mM NaH2PO4, 10 mM KCl, 1 mM MgSO4, 36 mM β-mercaptoethanol, 166 μl/ml lysozyme, 1.1 mg/ml ortho-nitrophenyl-β-galactoside (ONPG), and 6.7% v/v Fastbreak™ reagent) for enzyme activity measurements. β-galactosidase activity was monitored as a kinetic experiment at absorbance values of 420 nm(A_420nm_) and 595 nm (A595nm) on a Multiskan Spectrum microplate reader and Miller units were calculated using the modified Miller equation as described in (89) for samples tested in triplicate. For positive control transformants in the assay, 0.05 mM IPTG (final concentration) was added to media to induce P*tac lacZ* expression in pLacZ plasmids (Table 1) and compared to the same transformants in media lacking IPTG (negative control). Parental vector pME119EH served as the *lacZ* negative control to determine background assay absorbance values.

The high-throughput 96-well microplate β-galactosidase assay method used for broth cultures was modified to incorporate biofilm cultures as follows: MBEC devices used for AST did not produce sufficient quantities of biofilm cells (based on OD_600 nm_ values) for high-throughput 96-well microplate β-galactosidase assays. To produce sufficient biofilm cell titres for β-galactosidase assays, we assembled a 1 ml deep well pegged lid biofilm device. Deep well pegged lid biofilm devices were formed by inserting a sterile autoclaved semi-skirted flat-deck 0.5 ml 96 well PCR tube tray (Thermo Scientific, catalogue number AB1400) was inserted into the wells of an autoclavable polypropylene 1.2 ml 96-deepwell storage microplates (Axygen; P-DW-12-HC-S, VWR; 89230-176); biofilms formed on the surface of sterile inserted PCR tray. Starting biofilm cultures of the BW25113 strain transformed with pMS119EH, placZ (+/- 0.05 mM IPTG), pPq, pPc were grown as described for AST MBEC device experiments in the deep well devices in 0.6 ml using LB-AMP broth for 24 hrs with shaking at 155 rpm. These deep well biofilm devices provided a larger surface area for biofilm formation and increased total biofilm cell yields by 4-8 fold as compared to traditional MBEC devices. Biofilms for β-galactosidase assays were recovered from PCR tube ‘peg lids’ after rinsing twice in sterile PBS then sonicating for 10 minutes in 25°C water bath in 0.56 ml per well Miller Custom Mix solution (0.5M Na_2_HPO_4_, 1M NaH_2_PO_4_, 1M KCl, 0.1M MgSO_4_, 1M β-mercaptoethanol, 1X FastBreak™) for use as the starting cultures where immediately before kinetic absorbance measurement 40μl of Miller Assay solution (11.67 mg/ml lysozyme and 5.5 mg/ml ONPG) was added to each well for the high-throughput β-galactosidase assay procedure previously described involving planktonic broth cultures (89). These experiments were performed in triplicate and Student’s T-tests were used to determine statistically significantly differences from controls or samples and two-tailed *p*-values ≤ 0.05 were deemed significant.

## Acknowledgements

We wish to thank Branden Gregorchuk for helpful manuscript comments and Nadine Lewis from BioBasic Inc. for her assistance with gene synthesis optimization. Funding for this study was provided to DCB by a Manitoba Medical Service Foundation operating grant [8-2016-4] and an NSERC-Discovery Grant [RGPIN-2016-05891].

## Supplementary Material captions

**Table S1**. A summary of SMR protein sequences encoded by proteobacterial plasmids, including protein annotations, accession numbers, taxonomy and isolation sources.

**Table S2**. A summary of the 500 nucleotide upstream regions for each respective *qac* gene identified from the surveyed plasmids examined in this study. BLASTn search result summaries (as of February 2019) were included using proteobacteria (taxid:1224) and broad host range plasmid (taxid# 36549) GenBank database searches involving the SMR query sequences listed in black section headers.

**Table S3**. A summary of the 500 nucleotide upstream region corresponding to *sugE(p)* sequences identified from plasmid/integron sequences and BLASTn search results (as of February 2019) of broad host range plasmid (taxid# 36549) GenBank databases using the query amino acid sequence *E.coli sugE* YP_002302254 and *E.coli* gdx (NP_418572.4). Bolded sequences highlight *E. coli* and *P. aeruginosa gdx* genomic regions used as reference sequences for upstream homology.

**Figure S1**. COBALT alignment of 533 SMR protein sequences detected on proteobacterial plasmids. Sequence information corresponding to each COBALT sequence tag in the alignment ‘lcl’ is provided in Table S1.

**Figure S2**. The final multiple sequence alignment of the 20 unique Qac and SugE protein sequences. Boxed Qac residues highlight amino acid residues that vary when compared between other Qac sequences. Green boxes highlight conserved SugE residues. The alignment was generated in Jalview v2.10.5 [85].

**Figure S3**. 16% (T) SDS-Tricine PAGE analysis of isolated total cell membranes isolated from A) BW25113 and B) and KAM32 SMR transformants to determine SMR protein accumulation. Protein bands were imaged by UV irradiation using 0.5% (v/v) trichloroethanol added to the gels during casting [63] and loading dye was run off the bottom of the gel for 25 minutes to resolve bands shown in both panel images. Panel A shows total membrane protein extracts isolated from BW25113 transformants over expressing each cloned SMR in the study. The total membrane protein amounts loaded in each well was 25μg. Panel B shows total membrane protein extracts isolated from KAM32 transformants overexpressing each cloned SMR in the study. The total membrane protein loaded in each well of the gel in panel B was 25ug. In both panels, cultures grown for membrane extractions reached 0.5 units at OD_600nm_ and were induced to overexpress its respective SMR gene with 0.05 mM IPTG for 3 hrs at 37°C prior to harvesting cell pellets by centrifugation. Membrane isolations were performed as described by [63], and total membrane proteins concentrations were determined by Modified Lowry Assay. All membrane extract samples were standardized by diluting based on Lowry concentrations in nuclease free water before being mixed into sample loading buffer (100 mM dithiothreitol, 150 mM Tris-HCl pH 7, 12% (w/v) SDS, 30% (w/v) glycerol, 0.05% Coomassie Brilliant Blue G250 dye) at a 2:3 sample: loading dye volume ratio prior to loading in wells of gel.

**Figure S4**. Growth curves of BW25113 and KAM32 SMR transformants to determine optimal IPTG induction levels. In all panels, final concentrations of IPTG from 0-500 mM were individually added to wells containing standardized (OD_600nm_ 1.0) 10^−3^ diluted cultures in fresh LB-AMP and grown for 24 hrs at 37°C with shaking in 96 well microtiter plates. OD_600 nm_ values collected over 24 hrs measured in 30 min intervals is shown in each panel. Growth curves of BW25113 (A-D) or KAM32 (E-H) strains transformed with pMS119EH (A, E) pEmrE transformants (C, F), pQacE (C,G), and pGdx (D,H) exposed to various IPTG levels are shown. Trends observed for pGdx were also observed for pSugE(p).

**Figure S5**. Summary alignments of the upstream nt region (−115 to +1) corresponding to plasmid encoded *qacE, qacEΔ1, qacF, qacG*, and *qacH* genes examined in this study. Plasmids with 100% nt sequence identity in the – 115 to +1 region were omitted from the alignments shown in each panel to highlight unique sequences; all sequence information is summarized in Table S2. In panels A-C, the upstream *E.coli* (Ecol) *emrE* sequence is included at the top of each alignment as a reference and a consensus plot is provided below each alignment to highlight conserved nt identified at each aligned position. Pq promoter sequence identity is highlighted in yellow. A) Upstream regions of plasmid encoded *qacEΔ1* sequences. The −115 upstream region from *qacEΔ1* show 98-100% identity to the class I integron Pq promoter region. B) Upstream regions of plasmid encoded *qacF* and *qacH* genes. Few *qacF* sequences had identity to the Pq promoter (yellow). Pq promoter identity is highlighted in bold on the consensus plot below the *qacF/ qacH* alignment. C) Upstream regions of plasmid encoded *qacE* and *qacG* sequences. Upstream regions of *qacG* showed little identity to any SMR gene upstream regions. Species abbreviations: *Achromobacter denitrificans* (Aden), *Acinetobacter baumannii* (Abau), Citrobacter freundii (Cfre) *E. coli* (Ecol), *Enterobacter cloacae* (Eclo), *E. hormaechei* (Ehor), *Edwardsiella tarda* (Etar), *Klebsiella pneumoniae* (Kpne), *K. oxytoca* (Koxy), *K. variicola* (Kvar), *Leclercia adecarboxylata* (Lade), *Pseudomonas aeruginosa* (Paer), *P. alcaligenes* (Palc), *Salmonella enterica* (Sent), *Serratia marcescens* (Smar), *Stenotrophomonas maltophilia* (Smal).

**Figure S6**. Genetic arrangements of various SMR genes with respect to their promoter regions from various plasmids.

**Figure S7**. A summary of aligned putative 5’ UTR regions of plasmid encoded gdx/ sugE genes. In both panels, all identical sequences (100% sequence identity) within the ClustalO alignment of 115 plasmids with a *gdx/ sugE* sequence were omitted and only unique regions 65 nt upstream from the +1 start site of the gene are shown. Accession numbers listed on the right-hand side of each sequence corresponds to the specific plasmid each *sugE* gene was identified from. **A)** An alignment of the 5’ UTR upstream regions corresponding to plasmid encoded *sugE(p)* genes with 100% amino acid identity to YP_002302254.1. **B)** Aligned 5’UTR regions corresponding to *gdx/ sugE* sequence shown in Fig. 1A. In both panels, the coloured boxed nucleotides indicate high sequence identity (>99%) to the Gdm^+^ type-II riboswitch of *E. coli gdx* (NP.418572.4); putative P1 (red) and P2 (blue) Gdm^+^ type II riboswitch stem loops are indicated. Refer to Table S3 for plasmid accession number details.

**Figure S8**. Class I integron Pq and Pc promoters and riboswitch sequences studied. A) Diagram of a Class I integron and promoter regions with nucleotide sequences showing the commonly identified Pc and Pq promoter regions identified in this study. The Pc promoter is known to have a ‘weak’ (−35 TGGACA and −10 ‘TAAGCT’) and ‘strong’ (−35 TTGACA and −10 ‘TAAACT’) promoter version (38), where the more commonly identified weak promoter identified herein is shown. Uppercase single nucleotide letters indicate regions of identity between Pq and Pc promoter core sequences we determined in this study. B) A summary table of various Gdm^+^ riboswitch types, sequence accession numbers and sequences examined by BLASTn searches in this study.

**File S1**. Supplementary methods file describing promoter *lacZ* reporter vector cloning and primers used in the study.

